# Neuronal birthdate reveals topography in a vestibular brainstem circuit for gaze stabilization

**DOI:** 10.1101/2022.10.21.513243

**Authors:** Dena Goldblatt, Stephanie Huang, Marie R. Greaney, Kyla R. Hamling, Venkatakaushik Voleti, Citlali Perez-Campos, Kripa B. Patel, Wenze Li, Elizabeth M. C. Hillman, Martha W. Bagnall, David Schoppik

**Affiliations:** Depts. of Otolaryngology, Neuroscience & Physiology, and the Neuroscience Institute, New York University Grossman School of Medicine, New York, NY, 10016; Center for Neural Science, New York University, New York NY, 10004; University of Chicago, Chicago IL, 60637; Mortimer B. Zuckerman Mind Brain Behavior Institute, Columbia University, New York NY, 10027; Department of Neuroscience, Washington University, St. Louis MO, 63130

## Abstract

Across the nervous system, neurons with similar attributes are topographically organized. This topography reflects developmental pressures. Oddly, vestibular (balance) nuclei are thought to be disorganized. By measuring activity in birthdated neurons, we revealed a functional map within the central vestibular projection nucleus that stabilizes gaze in the larval zebrafish. We first discovered that both somatic position and stimulus selectivity follow projection neuron birthdate. Next, with electron microscopy and loss-of-function assays, we found that patterns of peripheral innervation to projection neurons were similarly organized by birthdate. Lastly, birthdate revealed spatial patterns of axonal arborization and synapse formation to projection neuron outputs. Collectively, we find that development reveals previously hidden organization to the input, processing, and output layers of a highly-conserved vertebrate sensorimotor circuit. The spatial and temporal attributes we uncover constrain the developmental mechanisms that may specify the fate, function, and organization of vestibulo-ocular reflex neurons. More broadly, our data suggest that, like invertebrates, temporal mechanisms may assemble vertebrate sensorimotor architecture.

## INTRODUCTION

Neurons are often organized according to shared functional properties. Such topography exists among sensory^1–4^ and motor^5–9^ nuclei, reflects developmental history^10–12^, and can facilitate computation^13–17^. In contrast, many brainstem nuclei appear locally disordered. One canonical example are the vestibular (balance) nuclei. While different vestibular nuclei can be coarsely distinguished by their projection targets^18–20^, individual nuclei appear largely disorganized^21–25^.

The apparent absence of topography is puzzling given the importance of sensory selectivity for vestibular circuit function. For example, the vertebrate vestibulo-ocular reflex uses a set of peripheral, central projection, and motor neurons (Figure 1A) to stabilize gaze after head/body movements^26–29^. Compensatory eye rotation behavior in the vertical axis requires stereotyped connectivity between two distinct sensory channels: one that responds exclusively to nose-up pitch-tilts and projects selectively to motor pools that rotate the eyes down, and one for the converse transformation (nose-down to eyes-up). However, no up/down organization among central vestibular projection neurons has been reported. Thus, similar to most brainstem nuclei, the organizational logic of neurons that comprise the vertical gaze stabilization circuit remains unresolved.

**Figure 1:**
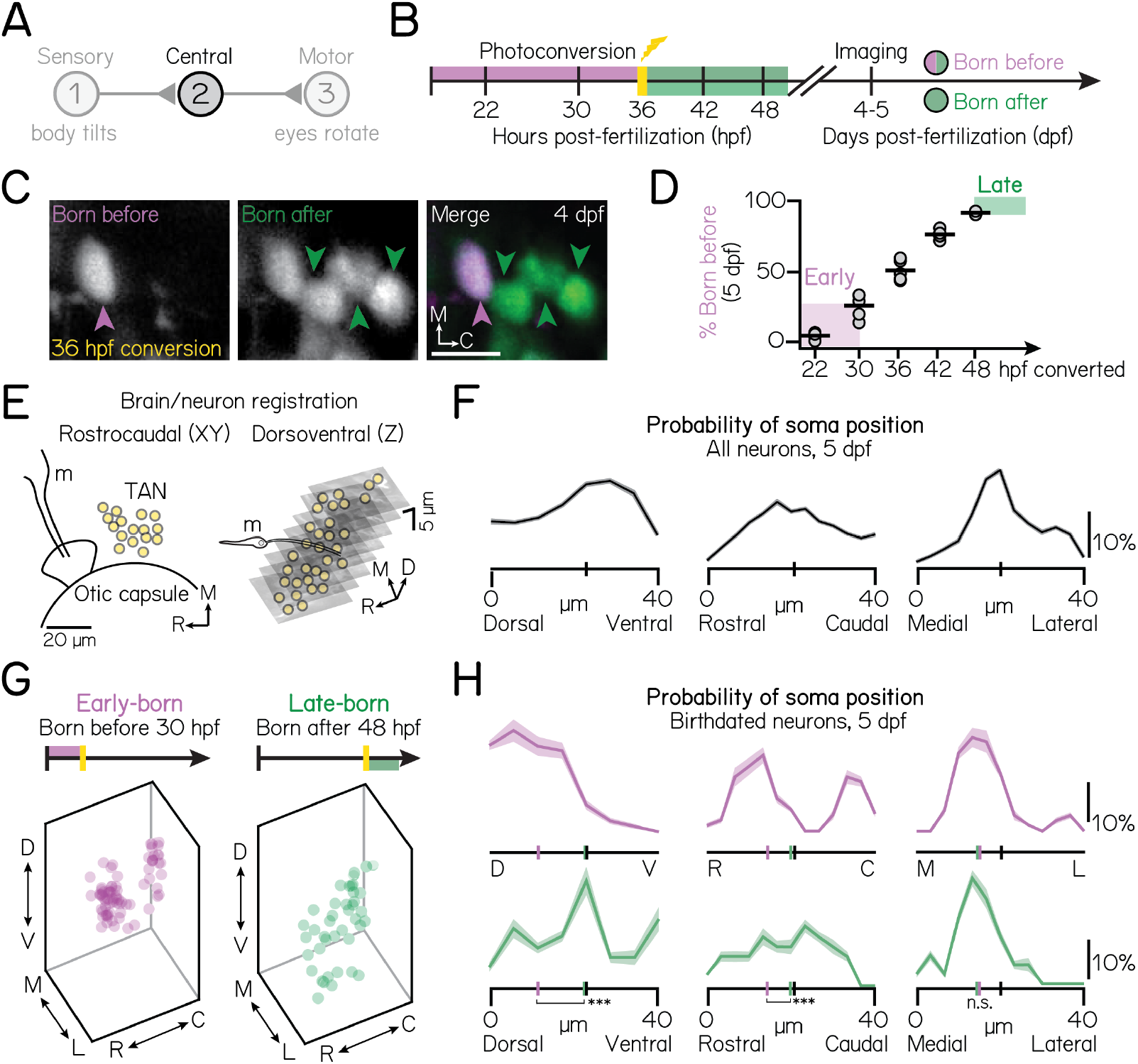
Projection neuron birthdate predicts soma position. (A) Schematic of the vestibulo-ocular reflex circuit used for gaze stabilization. (B) Timeline of birthdating experiments. Neurons born by the time of photoconversion have both magenta (converted) and green (unconverted) Kaede. Neurons born after photoconversion have exclusively green (unconverted) Kaede. All fish are imaged well after photoconversion, at 4-5 days post-fertilization (dpf). (C) Example projection neurons photoconverted at 36 hpf, visualized using the *Tg(-6.7Tru.Hcrtr2:GAL4-VP16; UAS-E1b:Kaede)* line. Magenta and green arrows indicate neurons born before or after 36 hpf, respectively. Scale bar (white), 20 μm. (D) Percent of projection neurons born at each timepoint. Black lines indicate the mean percent across individual fish (circles). Shaded bars indicate cohorts designated as early-born (converted by/born before 30 hpf) or late-born (unconverted at/born after 48 hpf). n=5 hemispheres/timepoint from at least N=3 larvae (22-42 hpf) or n=3 hemispheres from N=3 larvae (48 hpf). (E) Schematic of brain/neuron registration in the rostrocaudal/XY (left) and dorsoventral/Z (right) axes. Black solid lines outline anatomical landmarks used for registration. Yellow circles indicate the position of individual projection neurons. m, Mauthner neuron. (F) Soma positions of n=660 registered neurons from N = 10 non-birthdated fish at 5 dpf. Probability distributions shown are the mean (solid) and standard deviation (shaded outline) after jackknife resampling. Short vertical axis lines indicate the median position of all registered neurons. (G) Soma position of birthdated projection neurons. Early-born (left): n=74 neurons from N=7 fish. Late-born (right): n=41 neurons from N = 10 fish. (H) Same data as Figure 1G, shown as probability distributions for each spatial axis. Distributions shown are the mean (solid) and standard deviation (shaded outline) after jackknife resampling. Short vertical axis lines indicate the median position of early- (magenta) or late-born (green) projection neurons, compared to control distributions shown in Figure 1F (black). Three stars denotes a significant difference between the early- and late-born probability distributions at the p<0.001 level; one star, significance at p<0.05; n.s., not significant. See also: Figure S1.

Developmental time organizes somatic position, circuit architecture, and neuronal function. In *Drosophila* and *C. elegans*, temporal lineage underlies neuronal cell fate^30–32^, axonal connectivity^1,33–35^, and circuit membership^36,37^. In vertebrates, neuronal birthdate is correlated with the anatomical and functional properties of neurons^33,38–42^ and circuits^43–45^. These findings suggest that ontogeny might resolve outstanding mysteries of vestibular circuit organization. The bimodal sensitivity of vertical vestibulo-ocular reflex neurons makes them an excellent model, but in mammals, their early *in utero* development constrains exploring their emergent organization.

The larval zebrafish is uniquely well-poised to explore the development of the vestibulo-ocular reflex circuit^29^. Zebrafish are vertebrates that develop externally, are optically transparent^46–49^, and perform a vertical vestibulo-ocular reflex by 5 days post-fertilization^27,28,50^. Larvae possess both the semicircular canal and otolithic vestibular end-organs^27,51,52^, stimulation of which elicits reliable responses from central projection neurons^48,53,54^. Prior work established that vertical (eyes-up and eyes-down) motor neurons are spatially and developmentally organized^8^ and suggested that similar principles might organize the vestibular periphery^53,55^. Whether development organizes central projection neurons – and thus the vestibulo-ocular reflex circuit – remains unclear.

Here, we characterized the anatomy and sensory selectivity of optically-birthdated ^56^ projection neurons. By leveraging development, we discovered somatic topography among nose-up and nose-down projection neuron subtypes. Calcium imaging, electron microscopy, and loss-of-function assays established that development organizes the vestibulo-ocular reflex circuit, from pre-synaptic peripheral inputs to post-synaptic connectivity. We conclude that development reveals the “missing” topography in the vestibular system, challenging and extending existing models of the neural architecture for balance. This topography, and the developmental pressures it may reflect, offers considerable insights into the mechanisms that may specify the fate, function, and organization of vestibulo-ocular reflex neurons. More broadly, our data support a model in which temporal forces shape the assembly and function of a complete vertebrate sensorimotor circuit.

## RESULTS

### Central vestibular projection neurons are topographically organized by their birthdate

Our experiments required molecular access to the projection neurons that comprise a brainstem vestibular nucleus. We used a bipartite transgenic approach consisting of a Gal4 driver line, *Tg(-6.7Tru.Hcrtr2:GAL4-VP16)* and different UAS reporter lines (Methods) to access projection neurons. Projection neurons labeled in this driver line are located in the tangential vestibular nucleus, are collectively indispensable for the vertical vestibuloocular reflex, and can individually and collectively induce eye rotations^28,57^. To initially localize projection neuron somata, we used the *Tg(UAS-E1b:Kaede)* line^58^ to perform optical retrograde labeling^45^ of projection neuron axons at their ocular motor neuron targets (cranial nucleus nIII/nIV). Projection neuron somata were localized to a cluster at the lateral edge of rhombomeres 4-5, spanning approximately 40 μm in each spatial axis (Figures S1A to S1C). This cluster was anatomically bound ventrolaterally by the otic capsule, dorsomedially by the lateral longitudinal fasciculus, and rostrally by the Mauthner neuron lateral dendrite (Figures S1D to S1K). Somatic position and projection patterns are consistent with previous descriptions of the tangential vestibular nucleus^22,27,59–63^. On average, we observed 37±7 neurons per hemisphere (n=39 hemispheres, N = 30 larvae). Our approach permits reliable access to, and localization of, the projection neurons in the tangential vestibular nucleus responsible for the vertical vestibulo-ocular reflex.

To define when projection neurons develop, we optically labeled whole embryos, expressing *Tg(UAS-E1b:Kaede)*, at distinct, experimenter-defined timepoints (Figure 1B) ^56^. We imaged projection neurons in “birthdated” larvae at 5 days post-fertilization (dpf) using a confocal microscope (Figure 1C). We observed a linear increase in the number of optically-tagged projection neurons from 22-48 hours post-fertilization (hpf) (Figure 1D), at which point the majority (92% ±1%) of neurons were labeled (n=5 hemispheres/timepoint, 22-42 hpf; n=3 hemispheres, 48 hpf). Based on these data, we selected two temporally-defined populations for further comparative analyses: a cohort of early-born neurons born before (converted by) 30 hpf (25%±9% of projection neurons), and a late-born cohort of neurons born after (unconverted after) 48 hpf (8%±1%). Together, these populations comprise approximately one-third of projection neurons.

To determine if neuronal birthdate might reveal somatic topography, we manually registered all neurons to a common framework (Methods, Figures 1E and 1F). We discovered distinct, temporally-associated somatic organization (Figures 1G and 1H). Early-born somata were exclusively observed in the dorsomedial nucleus (n=74 neurons, N = 7 larvae), while late-born neurons were preferentially ventral (n=40 neurons, N=10 larvae). Spatial separation was significant across the extent of the nucleus (oneway multivariate analysis of variance, p=5.1*10^-14^), separately in the dorsoventral (two-tailed, two-sample KS test, p=6.17*10^-6^) and rostrocaudal (two-tailed, two-sample KS test, p=0.02) axes, and relative to chance (one-way multivariate analysis of variance, mean p=0.51±0.32). We observed no organization in the mediolateral axis (two-way, two-sample KS test, p=0.89). Our findings establish that neuronal birthdate predicts somatic position and reveals somatic topography within the tangential vestibular nucleus.

### Pitch-tilt stimuli differentiate two cardinal subtypes (nose-up/nose-down) of projection neurons

Projection neurons responsible for the vertical vestibulo-ocular reflex are directionally selective and respond primarily to either nose-up or nose-down head/body tilts (Figure 2A)^64–66^. To categorize the directional selectivity of individual projection neurons, we used Tilt In Place Microscopy (TIPM) with a two-photon microscope ^54^ (Figure 2B). Briefly, we used a galvanometer to rotate to, and hold larvae at an eccentric angle (either 19° nose-up or nose-down). We then rapidly (~4msec) rotated the galvanometer back to horizontal and measured the change in fluorescence of a genetically-encoded calcium indicator (UAS:GCaMP6s). As the duration of the step was orders of magnitude faster than the time constant of GCaMP6s^67^, the fluorescence on return to horizontal predominantly reflects neuronal activity at the previous eccentric position, and can reliably classify directional sensitivity in comparable projection neurons^54^.

**Figure 2:**
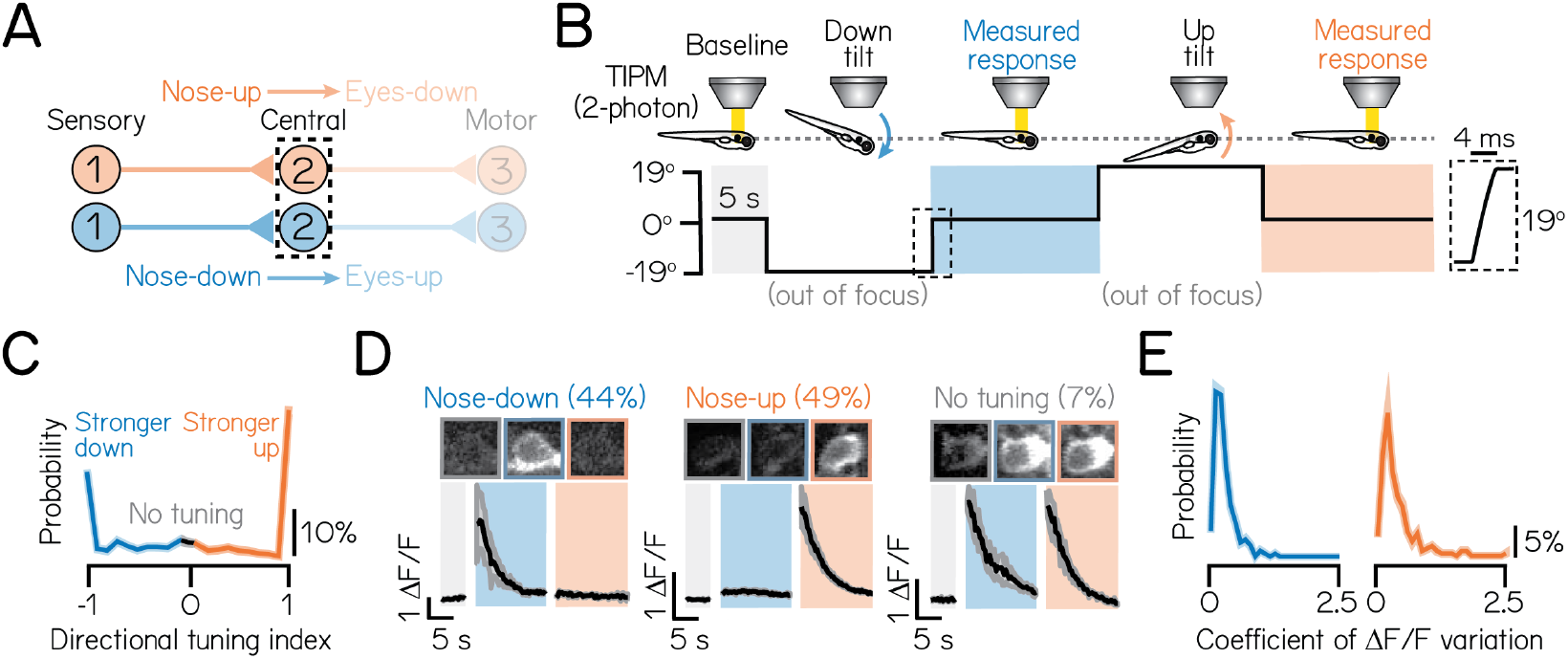
Pitch-tilt rotations reliably differentiate two cardinal subtypes of projection neurons. (A) Circuit schematic for pitch-tilt rotation experiments. Black dashed lines outline projection neurons as circuit population of focus. (B) Schematic of pitch-tilt rotation stimulus and imaging using Tilt In Place Microscopy (TIPM) and a two-photon microscope. Shaded regions show time of measured responses following nose-down or nose-down tilts. Inset shows a feedback trace from the galvonometer during the restoration step to horizontal. (C) Distribution of the directional tuning for all sampled neurons. Grey region indicates neurons with no directional tuning; blue and orange regions indicate neurons with stronger nose-down or nose-up responses, respectively. Criteria detailed in Methods. Solid line shows mean from jackknife resampling; shaded bars, standard deviation. (D) Example images and traces of a nose-down projection neuron (left), nose-up projection neuron (middle), and a projection neuron with no directional tuning (right) during TIPM. Projection neurons are visualized using the *Tg(-6.7Tru.Hcrtr2:GAL4-VP16)* line and express a UAS:GCaMP6s calcium indicator. Solid black lines show mean response across three stimulus repeats; shaded lines, standard deviation. Parentheses indicate the percent of neurons with each subtype, n=467 neurons, N=22 fish. (E) Distributions of the coefficient of variation of peak ΔF/F responses across three stimulus repeats for nose-down (left) and nose-up (right) neurons. Solid lines shows mean from jackknife resampling; shaded bars, standard deviation. See also: Figures S2 and S3.

We categorized the subtype identity (nose-up/nose-down) of 471 projection neurons from 22 larvae according to the direction(s) that evoked excitation. Approximately half of all neurons (46%) produced measurable excitation to both nose-up and nose-down tilts. To describe a neuron’s directional selectivity and assign a subtype identity, we compared the magnitude of the excitatory response to nose-up and nose-down tilts. We defined a directionality index from −1 (exclusive nose-down selectivity) to +1 (nose-up) and a selectivity threshold (>0.1, or ~22% difference in response magnitude; Methods). Most neurons (93%) were selective for only one direction (Figures 2C and 2D; nose-down mean: −0.73 ±0.30; nose-up mean: 0.84±0.28). We observed a nearly equal distribution of subtypes in our sample (44% nose-down, 49% nose-up, 7% no directional preference). Importantly, directional responses were consistent across repeated tilts (Figure 2E; mean coefficient of variation, nose-down: 0.27±0.21%; nose-up: 0.39±0.36%), with no variability in directional selectivity. TIPM can therefore be used to classify projection neuron subtypes.

We next used TIPM to characterize how projection neuron subtypes respond to tonic tilt stimuli. Projection neurons responded to non-preferred directions in one of three ways: 1) no response (Figures S2A and S2B), 2) weaker or equivalent excitation (Figure S2C), or 3) suppression, followed by above-baseline recovery (Figure S2D); classification detailed in Methods. Nose-down neurons produced stronger calcium responses to 19° pitch-tilts than nose-up neurons (one-tailed, two-sample KS test, p=6.5*10^-7^; mean nose-down ΔF/F: 1.97 ±1.65; mean nose-up ΔF/F: 1.21±1.10) and responded to non-preferred directions with weaker excitation (55% of nose-down neurons). Nose-up neurons primarily responded to non-preferred directional stimuli with suppression and above-baseline recovery (61% of nose-up neurons). This may reflect different sources of sensory input to nose-up and nose-down neurons or intrinsic differences in their encoding properties.

Vertical vestibular sensation is encoded by two different end organs in the inner ear: the utricular otoliths and the anterior/posterior semicircular canals. We adopted a loss-of-function approach to differentiate the contribution of each set of endorgans to tonic tilt responses. To assay the extent of the response that derived from the utricular otoliths, we presented pitch-tilt rotations to larvae with mutations in the *otogelin* gene^68^ (Figures S3A and S3B). *Otogelin* is expressed selectively in the inner ear^69^. Null *otogelin* mutants fail to develop utricular otoliths in the first two weeks, while heterozygous siblings are morphologically and functionally intact^70^. Projection neurons in *otogelin* mutants (n=183 neurons, N = 3 larvae) exhibited significantly weaker calcium responses to pitch-tilt rotations compared to sibling controls (Figure S3C; n = 144 neurons, N=3 larvae; one-tailed, two-sample KS test, p=3.6*10^-16^l; mean control ΔF/F: 1.31±1.21; mean mutant ΔF/F: 0.36±0.43). To assay the contribution of the semicircular canals, we performed acute, unilateral lesions of both the anterior (nose-up) and posterior (nose-down) canal branches of the VIII^th^ nerve (Figures S3D and S3E). We observed weaker pitch-tilt responses in projection neurons in lesioned hemispheres (n=49 neurons, n=4 larvae) compared to control hemispheres (Figure S3F; n = 57 neurons, n=4 larvae; one-tailed, two-sample KS test, p=5.0*10^-4^; mean control ΔF/F=0.88±0.63; mean lesioned ΔF/F=0.57±0.71). Thus, as expected, tonic pitch-tilt responses originate in the inner ear, predominantly from the utricular otoliths and secondarily from the semicircular canals.

### Birthdate predicts the functional organization of projection neurons

As birthdate anticipated soma position, we next asked if tilt sensitivity was similarly organized. To determine if birthdate-related topography predicted subtype identity, we performed TIPM on birthdated projection neurons (Figure 3A). Birth order predicted directional subtype preference at 5 dpf: early-born neurons almost exclusively adopted a nose-up subtype identity (n=74 neurons, N=10 larvae; 96% nose-up, 4% nose-down), while late-born neurons were predominantly nose-down (Figure 3A; n=40 neurons, N=10 larvae; 75% nose-down, 25% nose-up). To assay organization, we pseudocolored the same dataset of birthdated neurons (same data as shown in Figures 1G and 1H) according to their nose-up/nose-down directional preference. We observed a clear relationship between birthdate, somatic position, and subtype preference (Figure 3C).

**Figure 3:**
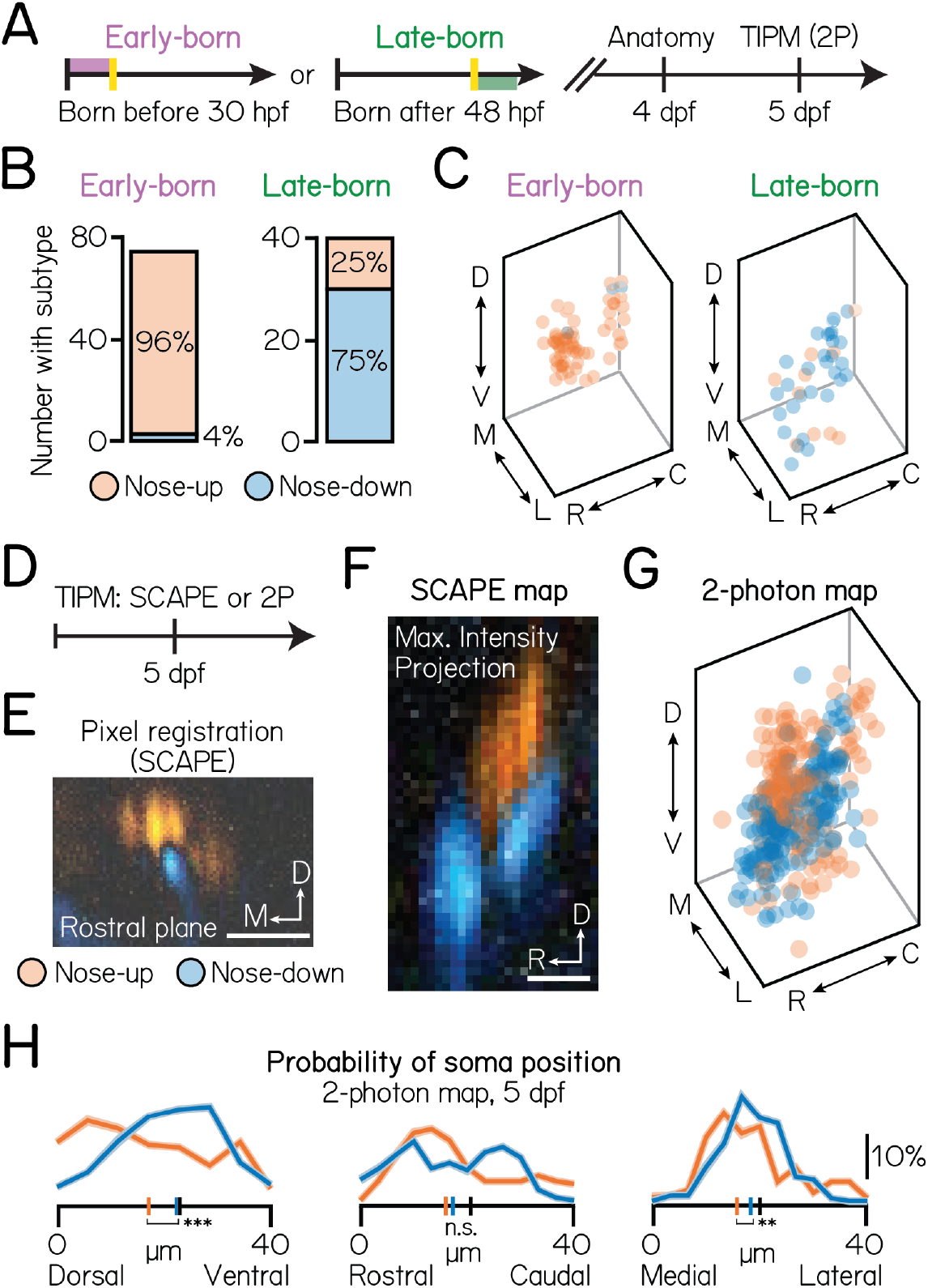
Birthdating reveals functional somatic topography to projection neurons. (A) Timeline of functional birthdating experiments. Larvae were birthdated at either 30 hpf or 48 hpf and then imaged on a confocal (4 dpf) and two-photon (5 dpf) to identify projection neuron soma position, birthdate, and directional (up/down) identity. Projection neurons were visualized using the *Tg(-6.7Tru.Hcrtr2:GAL4-VP16)* line and expressed both the UAS:Kaede and UAS:GCaMP6s indicators. (B) Number of early- (left) and late-born (right) projection neurons with each directional subtype identity. Data from same neurons shown in Figure 1. (C) Soma position of early- (left) and late-born (right) neurons, pseudocolored according to directional identity. Data from same neurons shown in Figure 1. (D) Timeline of topography validation experiments. TIPM was performed on non-birthdated larvae at 5 dpf as shown in Figure 2B, using either a SCAPE or two-photon microscope. Projection neurons were visualized using the *Tg(-6.7Tru.Hcrtr2:GAL4-VP16)* line and expressed only the UAS:GCaMP6s indicator. (E) Registration method for SCAPE experiments. Pixels are pseudocolored according to the direction that evoked a stronger response. One example rostral plane shown. (F) Maximum intensity projection of the entire tangential nucleus from one fish imaged with SCAPE, pseudocolored as described in Figure 3E. (G) Soma position of neurons imaged with two-photon TIPM, pseudocolored by directional identity. Soma registered using method shown in Figure 1E. Data from the same n=467, N=22 fish as in Figures 2C to 2E (H) Probability of soma position for nose-up (orange) and nose-down (blue) projection neurons from two-photon TIPM. Distributions shown are the mean (solid) and standard deviation (shaded outline) after jackknife resampling. Short vertical axis lines indicate the median position of up/down projection neurons, compared to control distributions shown in Figure 1F (black). Three stars denotes a significant difference between the nose-up and nose-down distributions at the p<0.001 level; two stars, significance at p<0.01; n.s., not significant. See also: Figure S4.

Early- and late-born cohorts comprise up to one-third of projection neurons. To test whether birthdated topography anticipates organization across the entire nucleus, we used a volumetric imaging approach. We performed TIPM using a swept, confocally-aligned planar excitation (SCAPE) microscope (Figures 3D and 3E)^49^. Subtypes were positioned into distinct, 10 μm clusters, complementing birthdated data (Figure 3F). Specifically, nose-up neurons occupied the dorsomedial-most extent of the nucleus, while nose-down neurons clustered into two ventrolateral stripes. Finally, we tested whether we could observe organization in our full (non-birthdated) two-photon dataset (Figures 3G and 3H). Spatial separation between subtypes was significant across the entire nucleus (n=360 mapped neurons, N=20 larvae; one-way multivariate analysis of variance, p=1.1*10^-5^), and separately in the dorsoventral (two-tailed, two-sample KS test, p=1.7*10^-6^) and mediolateral axes (two-tailed, two-sample KS test, p=0.01), and relative to chance (one-way multivariate analysis of variance, mean p=0.52±0.29, details of permutation testing in Methods). We also identified topography among neurons with similar non-preferred directional responses (Figure S4A) and response magnitudes (Figure S4B). As both of these features correlate with nose-up or nose-down selectivity, we propose that directional subtype is the simplest axis with which to study projection neuron topography. Taken together, our data links development with function to reveal unexpected topography within a single vestibular nucleus.

### Birthdate reveals structure to somatic assembly within the tangential nucleus

If birthdate predicts projection neuron topography, then longitudinal birthdating may reveal how this topography is assembled. We repeated TIPM birthdating experiments at intermediate timepoints (36 hpf and 42 hpf; Figures 4A and 4B). Given the small presence of late-born (after 48 hpf) nose-up neurons, we first tested whether subtypes are specified at a constant rate during nucleus differentiation or preferentially in a restricted temporal window. Competency for the nose-up subtype peaked at 36 hpf and then slowed as nose-down subtypes emerged (Figures 4C and 4D; 36 hpf: n=208 neurons, N = 7 fish; 42 hpf: n = 198 neurons, N = 5 fish; 30 hpf and 48 hpf, same data as Figures 3B and 3C). Nearly three-quarters of neurons born after 36 hpf adopted a nose-down fate (Figure 4E; 36 hpf: n=168 neurons, N = 7 fish; n=66; 42 hpf: n=66 neurons, N = 5 fish; 30 hpf and 48 hpf, same data as Figure Figures 3B and 3C). These patterns likely reflect opposing temporal gradients of cell fate specification that persist through nucleus differentiation (48 hpf), with a key transition point at 36 hpf.

**Figure 4:**
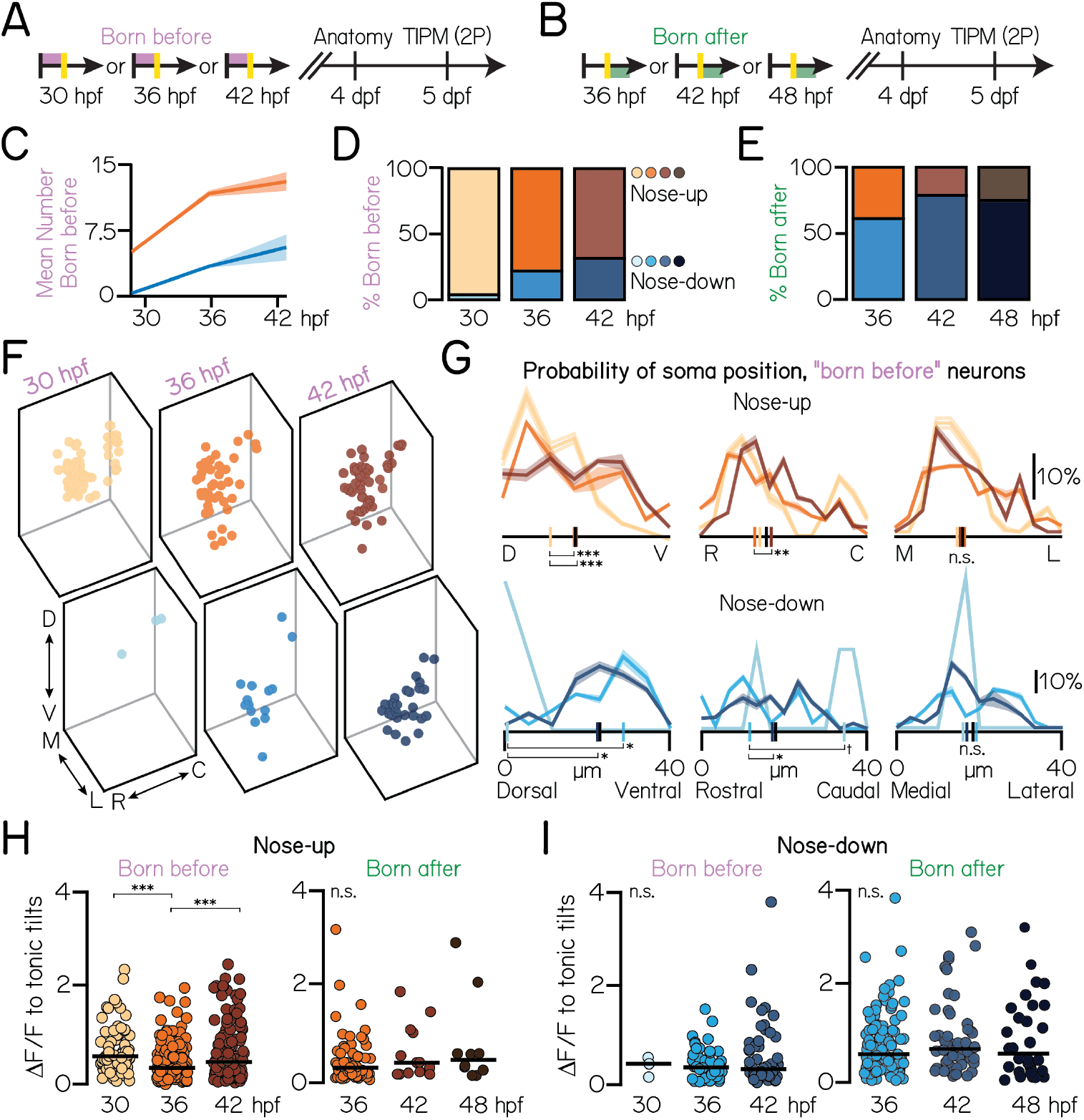
Projection neurons are assembled into the tangential nucleus in a stereotyped temporal sequence. (A) Timeline of functional birthdating experiments. Only projection neurons born before the time of conversion (red, converted Kaede) were analyzed. Projection neurons were visualized using the *Tg(-6.7Tru.Hcrtr2:GAL4-VP16)* line and expressed both the UAS:Kaede and UAS:GCaMP6s indicators. (B) Timeline of experiments. Only projection neurons born after the time of conversion (green, unconverted Kaede only) were analyzed. (C) Mean number of nose-up and nose-down neurons born before each age. Distributions shown are the mean (solid) and standard deviation (shaded outline) after jackknife resampling. (D) Percent of nose-up and nose-down neurons born before each age. Orange shades, nose-up; blue shades, nose-down. 30 hpf data is from the same N=7 fish as in Figures 1G, 1H, 3B and 3C. 36 hpf: n=208 sampled “born before” neurons from N=7 fish; 42 hpf: n=198 sampled “born before” neurons from N=5 fish. (E) Percent of sampled nose-up and nose-down neurons born after each photoconversion timepoint. Orange shades, nose-up; blue shades, nose-down. 36 hpf: n=168 “born after” neurons from N=7 fish; 42 hpf: n=66 “born after” neurons from n=5 fish. 48 hpf data same as in Figures 1G, 1H, 3B and 3C. (F) Soma position of neurons born before each age. 30 hpf: all data shown. 36 hpf: n=150/208 randomly selected neurons shown. 42 hpf: n=150/198 randomly selected neurons shown. (G) Probability of soma position for all “born before” neurons. Distributions shown are the mean (solid) and standard deviation (shaded outline) after jackknife resampling. Short vertical axis lines indicate the median position of all birthdated neurons, compared to control distributions shown in Figure 1F (black). (H) Maximum change in calcium fluorescence to tonic tilts for nose-up neurons born before (left) or after (right) each age. Each circle represents a unique neuron. Solid line shows the mean across neurons. (I) Maximum change in calcium fluorescence to tonic tilts for nose-down neurons born before or after each age. All data: Three stars denotes a significant difference at the p<0.001 level; two stars, significance at p<0.01; †, significance between p=0.08 and p=0.05; n.s., not significant.

We next measured how nose-up and nose-down neurons accumulated within the nucleus over time (Figures 4F and 4G). Nose-up neurons emerged from dorsal to ventral (Kruskal-Wallis test, p=4.1×10^-4^) and rostral to caudal (Kruskal-Wallis test, p=0.003). Nose-up topography was indistinguishable from 5 dpf control distributions (Figure 3H) by 36 hpf in the dorsoventral axis (Kruskal-Wallis test with multiple comparisons, p=0.002, 30 hpf vs. control; p=0.98, 36 hpf vs. control) and 42 hpf in the rostrocaudal axis (Kruskal-Wallis test with multiple comparisons, p=0.06, 36 hpf vs. control; p=0.63, 42 hpf vs. control). There was no temporal structure in the mediolateral axis (Kruskal-Wallis test with multiple comparisons, p=0.39). The two rostrocaudal “stripes” of nose-down neurons similarly emerged from dorso-rostral to ventro-caudal (dorsoventral: Kruskal-Wallis test, p=0.003; rostrocaudal: Kruskal-Wallis test, p=0.03). These “stripes” continued to develop through 36 hpf in the dorsoventral axis and 42 hpf in the rostrocaudal axis, at which point their distributions matched control (dorsoventral: Kruskal-Wallis test with multiple comparisons, p=0.05, 30 hpf vs. control; p=0.20, 36 hpf vs. control; rostrocaudal: Kruskal-Wallis test with multiple comparisons, p=0.07, 36 hpf vs. control; p=0.99, 36 hpf vs. control). This suggests that projection neuron topography is shaped early in development and contemporaneously with subtype specification.

We considered that late-born neurons of a particular subtype (up/down) may represent distinct sub-populations. To test this, we measured how strongly birthdated projection neurons responded to tilt sensation. We observed a small decrease in nose-up response strength at 36 hpf (Kruskal-Wallis test, p=1.8×10^-5^; Cohen’s d, 0.65, 30-36 hpf, −0.44, 36-42 hpf), but no difference in the distributions of late-born (“born after”) nose-up neurons (Figure 4H). We found no developmental trends for nose-down neurons (Kruskal-Wallis test, p=0.93; Figure 4I) or to other measurements, such as directional selectivity strength. We conclude that early- and late-born neurons of a particular subtype are functionally homogeneous. Thus, finer aspects of projection neuron function, except where linked to subtype (e.g., non-preferred response type), may be determined by non-temporal forces.

### Responses to high-frequency sensory stimulation are topographically organized by birthdate

Given the organization of projection neurons by tonic tilt responses, we predicted that their responses to other forms of vestibular end-organ sensory stimulation – and by extension, their VIII^th^ nerve sensory inputs – may be similarly organized. Projection neurons must also encode phasic (i.e. rapid) posture changes. Phasic changes to neuronal activity can be elicitepd with sufficiently high-frequency stimulation^54,71^, potentially following semicircular canal input to the tangential nucleus^22,25,72^. While their responses were weaker than to tonic tilts (Figure 5B), ~60% of neurons responded to impulses. Typically, neurons did not exhibit a clear preference for either impulse (Figure 5C; mean directional strength=0.06±0.39) with some variability in response magnitude across stimulus repeats (mean coefficient of variation: 0.85±0.41).

**Figure 5:**
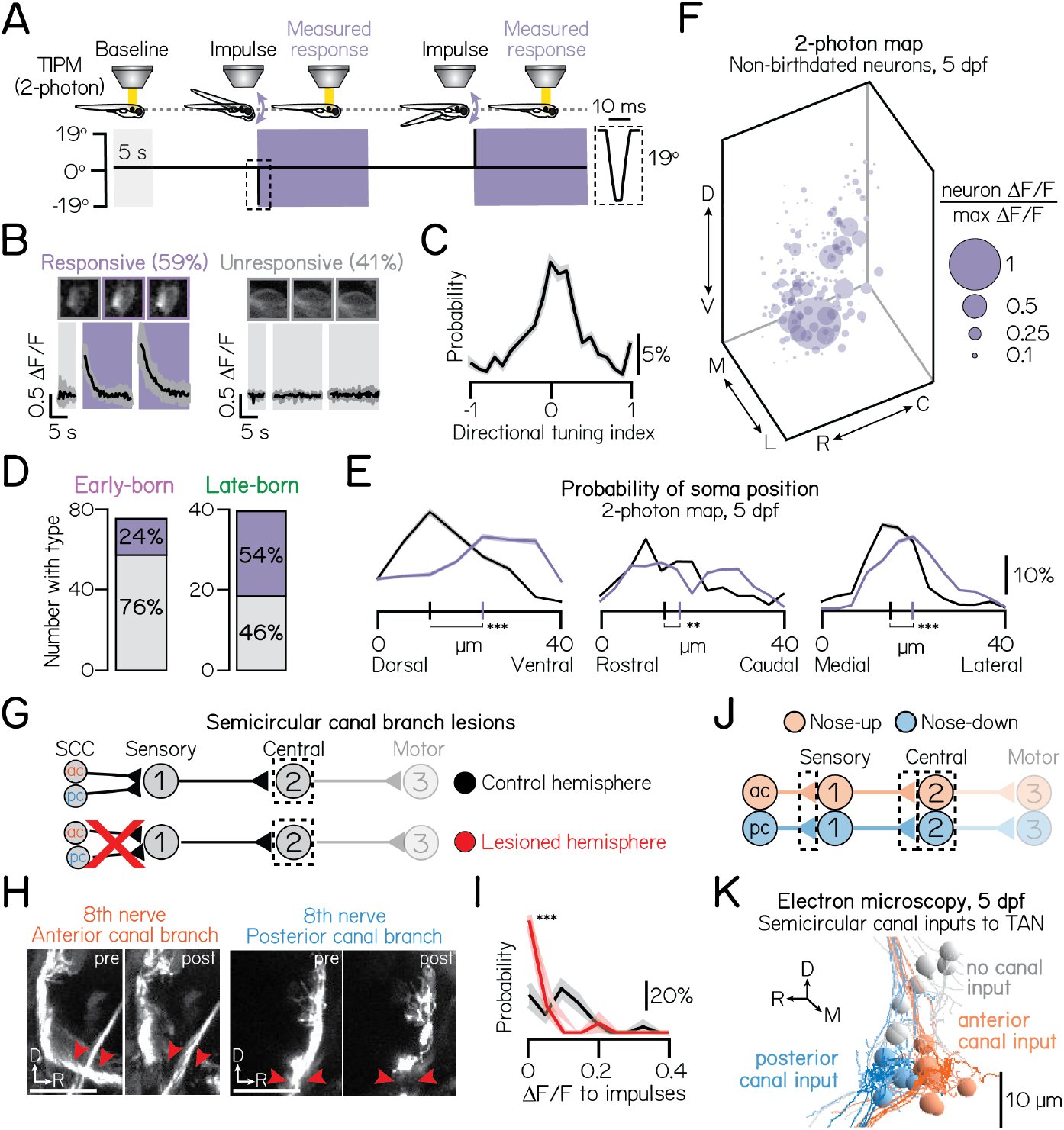
Birthdate reveals topography to semicircular canal-mediated, high-frequency stimulation responses. (A) Impulse stimulus waveform. Shaded bars indicate time in the horizontal calcium imaging plane. Dotted box shows zoom of 10 ms impulse. (B) Example traces from an impulse-responsive (left) and unresponsive (right) projection neuron. Parentheses show percent of neurons in sample. Same n=467 neurons, N=22 fish as Figure 2D. (C) Probability of directional selectivity to impulses. Zero indicates no directional preference. Distribution shows mean (solid line) and standard deviation (shaded outline) from jackknife resampling. (D) Number of early- and late-born projection neurons with (purple) or without (grey) impulse responses (purple). Data from same fish as in Figures 1G and 1H. (E) Probability of soma position for impulse-responsive (purple) or unresponsive (grey) neurons from non-birthdated, two-photon TIPM; same n=467 neurons, N=22 fish as Figure 2D. Short vertical axis lines indicate the median position. (F) Soma position of impulse-responsive neurons, scaled by impulse response strength (ΔF/F) relative to the strongest response observed. Larger circles indicate stronger responses. Data from the same neurons shown in Figure 3G. (G) Circuit schematic. Both the anterior and posterior semicircular canal branches of the VIII^th^ nerve are uni-laterally lesioned. Calcium responses of projection neurons (black dashed box) in lesioned and control hemispheres are compared. (H) Example images from larvae before and after uni-lateral VIII^th^ nerve lesions. Left and right image sets (replicated from Figure S3E) show the anterior and posterior semicircular canal branches, respectively, before and after lesion. Both branches are lesioned in each experiment. Red arrows point to lesion sites. VIII^th^ nerve visualized using the *Tg(-17.6isl2b:GFP)* line; larvae also expressed *Tg(-6.7Tru.Hcrtr2:GAL4-VP16; UAS:GCaMP6s* for projection neuron calcium imaging. (I) Probability distributions of the maximum ΔF/F response to impulse rotations in lesioned (red) and control (black) hemispheres. Responses shown only for the most ventral 15 μm of projection neurons. Solid lines shows mean from jackknife resampling; shaded bars, standard deviation. Control: n=68 impulse responses, N=3 fish. Mutants: n=132 impulse responses, N=3 fish. (J) Circuit schematic for electron microscopy experiments. Black dashed lines indicate the circuit elements of focus: synaptic connections from the anterior and posterior semicircular canals to first-order sensory neurons, synaptic connections from sensory neurons to projection neurons, and projection neurons. (K) Electron microscopy reconstruction of 19 projection neurons at 5 dpf. Soma pseudocolored based on innervation from sensory neurons that receive anterior semicircular canal input (orange) or posterior semicircular canal input (blue). Grey soma receive no semicircular canal input. All panels: Three stars denotes a significant difference at the p<0.001 level; two stars, significance at p<0.01. See also: Table S1.

Next, we assayed if the somata of impulse-responsive projection neurons were organized in space and time. We presented impulse stimuli to the same birthdated neurons described previously (n=74 early-born neurons from N = 7 larvae, n=40 late-born neurons from N = 10 larvae; same data as shown in Figure Figures 1G and 3C). We observed a notable relationship between neuronal birth order and impulse responsiveness: late-born neurons were twice as likely to respond to impulses (Figure 5D; 54% responsive) than early-born (24% responsive). As late-born neurons are ventrally localized (Figure 1H), we predicted that impulse responses may be similarly localized. Indeed, impulse responsive neurons were restricted to the ventral (two-tailed, two-sample KS test, p=22.7*10^-9^), lateral (two-tailed, two-sample KS test, p=5.9*10^-7^), and rostral (two-tailed, two-sample KS test, p=0.009) tangential vestibular nucleus (Figures 5E and 5F). Spatial separation was further significant across the entire nucleus (one-way multivariate analysis of variance, p=1.3*10^-9^) and relative to chance (one-way multivariate analysis of variance, mean p=0.46±0.30). We conclude that impulse-responsive projection neurons are born later and topographically constrained to the ventrolateral tangential vestibular nucleus.

### Organized sensory input from semicircular canal afferents underlies the topography of high-frequency stimulation responses

Given the ventral topography of impulse responses, we predicted that their VIII^th^ nerve sensory inputs (semicircular canals and/or utricular otoliths) may be similarly organized. To test whether the semicircular canals mediate impulse responses, we repeated our loss-of-function approach (Figures 5G and 5H). We measured responses in the ventral-most 15μm of projection neurons, which are maximally impulse-responsive (Figure 5F). Impulse responses in ventral neurons were almost entirely eliminated following canal lesions (one-tailed, two-sample KS test, p=1.1*10^-7^; mean control ΔF/F = 0.10±0.08; mean lesion ΔF/F = 0.02±0.05; control: n=24 impulse responses, N=4 larvae; lesion: n=16 impulse responses, N=4 larvae; Figure 5I). This effect was consistent across the entire nucleus (one-tailed, two-sample KS test, p=2.8*10^-4^; mean control ΔF/F: 0.08±0.07; mean lesion ΔF/F: 0.03±0.05; control: n=40 impulse responses, N=4 larvae; lesion: n=36 impulse responses, N=4 larvae).

To test whether impulse stimuli activate the utricular otoliths, we presented impulse stimuli to *otogelin* null larvae. Impulse responses decreased, but were not entirely eliminated, in the same ventral population on the *otogelin* background (one-tailed, two-sample KS test, p=1.7*10^-7^; mean control ΔF/F: 0.38±0.22; mean mutant ΔF/F: 0.14±0.10; control: n=43 impulse responses, N = 3 larvae; mutant: n=33 impulse responses, N = 3 larvae). We observed a similar effect across the entire nucleus (one-tailed, two-sample KS test, p=3.0*10^-22^; mean control ΔF/F: 0.25±0.22; mean mutant ΔF/F: 0.05±0.08; control: n=169 impulse responses, N = 3 larvae; mutant: n=129 impulse responses, N = 3 larvae). Together, these loss-of-function experiments support a primary role for the semicircular canals in mediating impulse stimuli responses, with a nominal additional contribution from the utricular otoliths.

We next adopted a connectomic approach to assay whether the observed impulse sensitivity of ventral neurons might reflect targeted inputs from anterior (nose-up) and posterior (nose-down) semicircular canal afferents. Using an existing set of serial electron micrographs^55^, we traced afferents from 5 dpf anterior (n=6 afferents) and posterior (n=5 afferents) semicircular canal cristae across two synaptic connections: to the statoacoustic ganglion, and then to projection neurons in the tangential vestibular nucleus (n=19 neurons; Figure 5J). We then pseudocolored projection neurons according to whether they received anterior, posterior, or no cristae input (Figure 5K). 12/19 identifiable projection neurons in the tangential nucleus (Methods, Electron Microscopy) received canal input, consistent with a previous anatomical report^22^. Input from the anterior and posterior canals was tightly organized along the rostrocaudal axis, matching the rostrocaudal organization previously reported in the VIII^th^ nerve^55^. Importantly, all canal-innervated projection neurons were located in the ventral-most 20 μm of the tangential vestibular nucleus.

We reasoned that projection neurons should receive directionally-matched input from both canal and utricular afferents. Utricular afferent directionality is derived from the orientation of their target hair cells in the macula. We traced 17 utricular afferents from their macular hair cell connections to projection neurons (Table S1) and predicted the directional tuning of these afferents based on the anatomical orientation of their hair cell inputs^55^. The vertical sensitivity of utricular and semicircular canal afferents matched perfectly in all canal-innervated projection neurons. Together, we conclude that semicircular canal input to projection neurons is tightly organized in space, matched to utricular afferent directionality, and preferentially targeted to later-born (ventral), impulse-responsive projection neurons.

### Axonal trajectories and synaptic connectivity to extraocular motor neurons follow development

Somatic topography, afferent input, and response properties are organized according to birthdate. We reasoned that birthdate might similarly predict spatial connectivity to downstream extraocular motor neuron targets. Previously, we identified spatiotemporal organization among the pools of extraocular motor neurons, located in cranial nuclei nIII and nIV, that control vertical/torsional eye movements. Motor neurons in cranial nucleus nIII innervate eyes-down muscles, are exclusively located dorsally, and born earlier than their eyes-up counterparts^8^. We predicted, given the shared spatial and temporal organization of motor neurons and projection neurons, that early-born projection neurons may preferentially target early-born motor neurons of the same temporal (early-born), spatial (dorsal), and functional (nose-up/eyes down) type.

To measure axonal organization, we birthdated projection neurons early in development (36 hpf) and then assayed the spatial extent of their axon trajectories at 3 dpf (Figures 6A and 6B). In all larvae (N = 3), axons from early-born (converted) somata projected exclusively along the dorsal medial longitudinal fasciculus. Photoconverted axons also contained a characteristic branch to dorsal extraocular motor neurons in nIV, showing they originate from the nose-up/eyes-down projection neuron subtype (Figure 6C)^28^. We conclude that the axons of early-born (nose-up) projection neurons project dorsally in the medial longitudinal fasciculus, targeting complementary early-born, dorsal motor neurons that move the eyes down.

**Figure 6:**
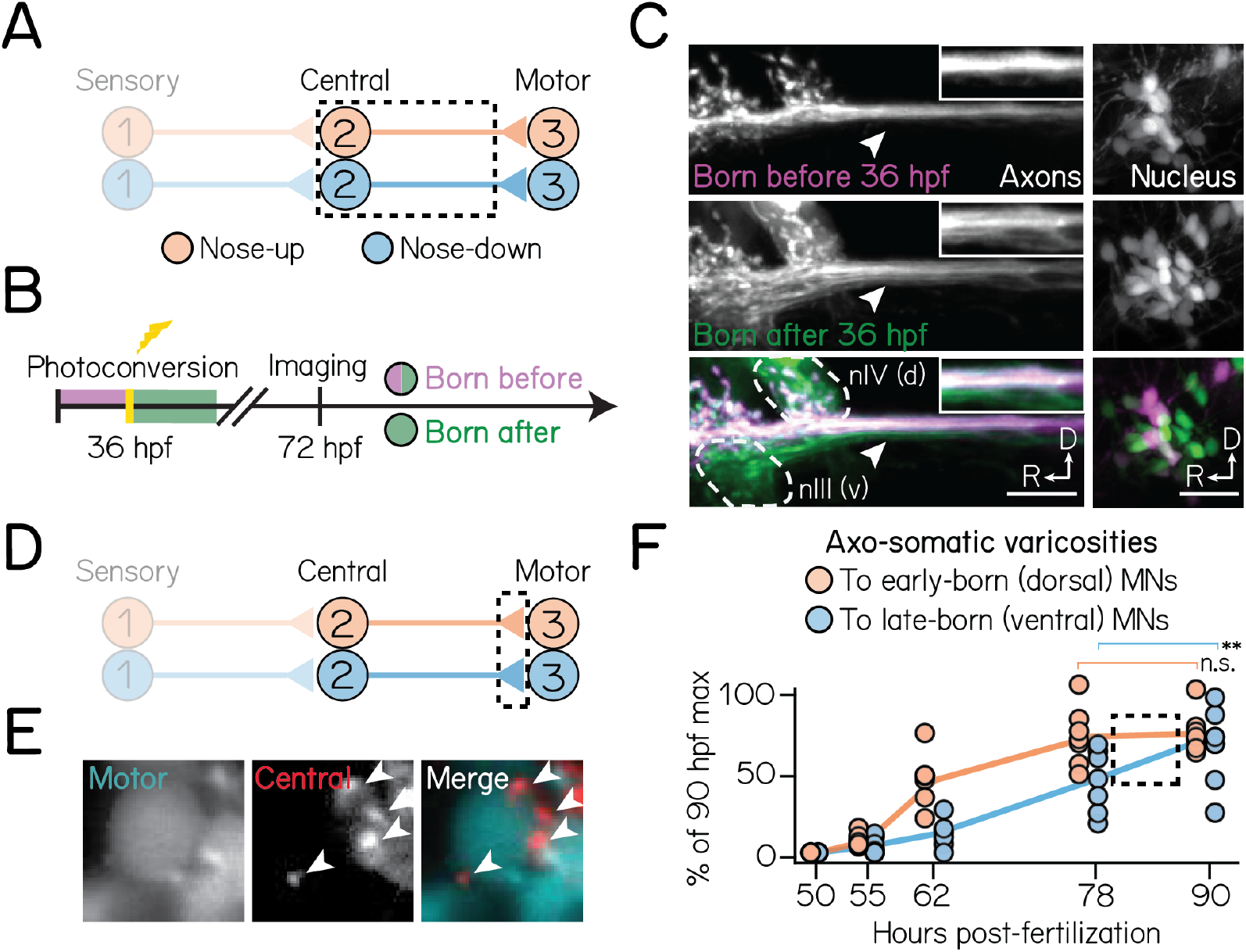
Birthdate predicts patterns of axonal trajectories and synapse formation between projection neurons and extraocular motor neurons. (A) Circuit schematic for axon birthdating experiments. Black dashed lines outline projection neuron soma and axonal projections to the extraocular motor nuclei. (B) Timeline of axon birthdating experiments. Larvae are only photoconverted at 36 hpf. (C) Birthdated axons from one example fish. Top row shows axons (left) from soma (right) born by 3 6 hpf; middle row shows axons that were not born by 36 hpf; bottom row shows merge. White arrows point to the ventral axon bundle. Inset shows zoom of axons. Dotted lines outline extraocular motor neuron somata. Scale bars, 20 μm. (D) Circuit schematic for synapse formation measurements. Black dashed lines outline axo-somatic varicosities from projection neurons onto post-synaptic ocular motor neurons. (E) Example image of an extraocular motor neuron (left) and axo-somatic varicosities (middle); right image shows merge. White arrows point to four example varicosities. Scale bar, 5 μm (F) Rate of varicosity growth to early- and late-born ocular motor neurons, quantified as as a percentage of the maximum number of varicosities observed at 90 hpf. Circles represent individual fish. Dotted box highlights the time where the rate of varicosity formation to motor neuron subtypes diverges. n ≥ 5 fish per timepoint. Two stars denotes a significant difference at the p<0.01 level; n.s., not significant.

We then tested whether projection neuron birthdate correlates with their synaptic development to eyes-up or eyes-down motor neurons. Vestibular projection neuron axons form varicosities onto motor neuron somata; varicosities co-localize with synaptic markers^73^. To assay synaptic development, we measured the number of axo-somatic varicosities from projection neurons onto either dorsal (early-born) or ventral (late-born) motor neuron somata (Figures 6D and 6E). Varicosities developed in three phases: initial onset (52-55 hpf), linear growth (55-78 hpf), and plateau (> 78 hpf) (Figure Figure 6F). The initial appearance of varicosities was comparable between all dorsal and ventral motor neurons. Varicosities to dorsal motor neurons developed rapidly, reaching their expected numbers (maximum observed at 90 hpf) by 78 hpf (one-way ANOVA with multiple comparisons, p=0.002, 62 hpf vs. 90 hpf; p=0.99, 78 hpf vs. 90 hpf). In contrast, varicosities to ventral motor neurons continued accumulating through 90 hpf (one-way ANOVA with multiple comparisons, p=0.01, 78 hpf vs. 90 hpf). This suggests that varicosities develop in a complementary dorsal-to-ventral pattern as motor neuron and projection neuron soma. Collectively, these findings reveal that projection neuron birthdate anticipates both axonal and synaptic assembly onto circuit-appropriate motor neuron outputs.

## DISCUSSION

Here, we show how previously hidden functional topography in the vestibular system emerges over development. Neuronal birthdating first uncovered spatial segregation between early- and late-born projection neurons. 2-photon and volumetric imaging with TIPM showed that directional selectivity to body tilts followed soma position and birthdate. We then combined loss-of-function and electron microscopy assays to show birthdate-related organization to upstream semicircular canal inputs. Lastly, we established that birthdate differentiates the timecourse of axon targeting and synapse development onto downstream ocular motor neurons. Taken together, we find that development reveals organization within the vestibular brainstem, its peripheral inputs, and its motor outputs (Figure 7). We propose that time plays a causal role in fate determination, topographic organization, and, by extension, vestibular circuit formation. Our data suggests mechanisms for projection neuron fate specification. More broadly, our findings offer insights into how time shapes vertebrate sensorimotor circuits.

**Figure 7:**
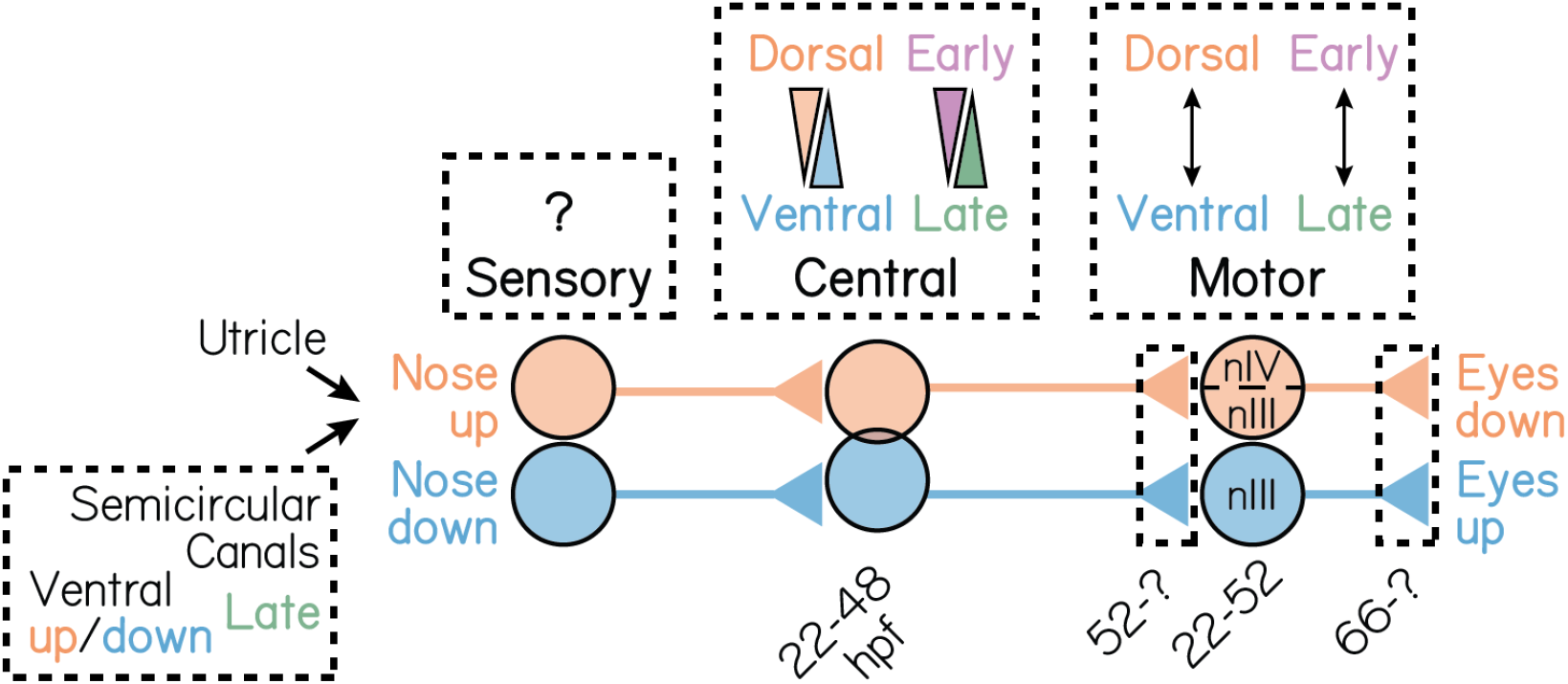
A model for the spatiotemporal organization of the gaze stabilization circuit. Spatiotemporal development of neurons that mediate the nose-up/eyes-down (orange) and nose-down/eyes-up (blue) gaze stabilization reflexes. nIII refers to the oculomotor cranial nucleus; nIV to the trochlear cranial nucleus IV.

### The impact of discovering spatiotemporal topography in the vestibular system

Our discovery of spatiotemporal topography relied on bringing new tools to bear on the classic question of vestibular circuit organization. Technological improvements have a long history of revealing organization in other sensory systems, from precise olfactory maps^74,75^ to the functional architecture of the visual system^76,77^. Prior characterizations had proposed that vestibular brainstem nuclei are disorganized^18–20,55,78^. Instead, retrograde tracing studies across chick, zebrafish, frog, and mouse had collectively identified coarse axonal projections – e.g., to spinal or ocular motor circuits – as their primary organizational axis^18–20,62^. Here, we used a modern toolkit consisting of: (1) a transgenic line of zebrafish that specifically labels vestibulo-ocular reflex projection neurons^28^, (2) *in vivo* birthdating^56^ early in development, and (3) spatial mapping of tilt responses^54^. These tools allowed us to extend the “axonal projection” model of vestibular brainstem organization to functional channels within an individual vestibular nucleus. More broadly, our results suggest that the vestibular brainstem is organized commonly to its sensory counterparts.

Topographic maps are the scaffold on which connectivity and function are built. Consequently, other model systems have used maps to illuminate the developmental origins of neuronal fate^79^, circuit assembly^35–37,80^, and function^42,81,82^. Relative to other “mapped” sensory systems, almost nothing is known about vestibular reflex circuit development^83^. Below, we discuss how our spatiotemporal maps illuminate how vestibulo-ocular projection neuron fate may be determined. The common temporal topography across channels of the vestibulo-ocular reflex circuit may similarly constrain its assembly. Finally, our data illustrate how principles of invertebrate circuit development in time may be implemented in vertebrate counterparts.

### The importance of birthdate in specifying vestibular projection neuron fate

Fate decisions are often the first step in constructing a neural circuit and occur upstream of the molecular logic that dictates circuit connectivity, functional attributes, and behavior^35,79,84–86^. What mechanisms might specify binary fate (nose-up/nose-down) in vestibular projection neurons? The absence of clearly delineated boundaries between nose-up and nose-down neurons argues against a role for spatial patterning programs. Projection neuron topography is neither laminar like cortical nuclei^87,88^ nor cleanly differentiated in one spatial axis like motor nuclei ^5,7–9,89,90^ or brainstem nuclei^18–20,91,92^. Although reaction-diffusion processes can produce periodic patterns in tissue^93,94^, the structure we see is, to the best of our knowledge, incompatible with known spatial cues. Across the hindbrain, broad spatial patterning signals act as the primary architects of rhombomeric topography^19,62,78,95–97^. We propose that *within* individual vestibular nuclei, the mechanisms that specify projection neuron fate lie elsewhere.

Work in comparable systems offers several additional models for binary fate choice^98^, many of which are inconsistent with our data. Specifically, our data argue against models with postmitotic implementation of stochastic choice, sister cell interactions, and lateral inhibition. One hallmark of a stochastic process is coincident appearance of subtypes, as in *Drosophila* R7 photoreceptors^99,100^. Similarly, cells with differential fates may be the result of interactions between coincidentally-born sister cells, as seen in v2a/v2b spinal interneurons^101,102^. Instead, we observe a clear temporal pattern to fate specification: early-born neurons are predominantly nose-up, while late-born neurons are nose-down. We can similarly eliminate a role for mutually-repressive (lateral inhibition) programs, in which local interactions between post-mitotic neurons specify fates. Such interactions establish patterned boundaries between subtypes^103^ and an unequal number of neurons in each subtype^104^. We observe that nose-up and nose-down neurons are roughly clustered and present in a 1:1 ratio. Thus, although binary (nose-up/nose-down) projection neuron fates are reminiscent of post-mitotic fate decisions in other sensory systems, they do not employ the same fate specification strategies.

Instead, projection neuron fate appears to be specified sequentially. Key evidence comes from their opposing rates of fate specification (Figure S4), where the appearance of nose-up neurons is greatest early in development and declines as nose-down probability rises. Such a competency “switch” must reflect differential molecular logic to specify fate. In the mouse optokinetic reflex circuit, directionally-selective (up/down) retinal ganglion cells employ Tbx5 in such a manner^105^. The similarities between the vestibulo-ocular reflex and optokinetic reflex circuits argue that a comparable process may instantiate vestibular projection neuron fates. Further evidence comes from the role of Fezf1 in the fate specification of on and off starburst amacrine cells in the retina^79^. Testing this hypothesis requires uncovering the transcriptional determinants of projection neuron fate. The clear temporal delineation between nose-up and nose-down channels, together with recent transcriptional analyses of vestibular neurons^106^, the zebrafish hindbrain^107,108^ and lineage tracing tools^109–111^, offer a basis for differential analyses of closely-related subtypes^79,105,112^. Looking ahead, defining the factors that instantiate projection neuron fate will be key to understand the assembly of the vestibulo-ocular reflex circuit.

### Circuit assembly: first-come, first served, or firstborn, first served?

The relationship between time and fate that we observe argues against the current “reverse order” model of vestibulo-ocular reflex circuit formation. Retrograde tracing experiments in the developing chick had suggested that projection neurons do not synapse onto extraocular motor neurons until the motor neurons connect to muscles^18^. This temporal delay led to the proposal that, for the vestibulo-ocular reflex circuit, “synaptic connectivity is established in reverse order to the signaling direction,” that is, from motor neuron on to muscle, followed by vestibular projection neuron on to motor neuron, and finally vestibular afferent on to vestibular projection neuron^83,113^. Instead, we find that vestibular projection neurons synapse on to extraocular motor neurons (Figure 6) well before those motor neurons synapse on to muscles^114^. Comparative approaches^115–117^ could resolve the question of whether the temporal differences we observe are zebrafish-specific. Together, our work rejects the current model for vestibulo-ocular reflex ontogeny.

Sensorimotor transformations require precise connectivity, and connectivity itself could therefore determine circuit assembly. For vestibulo-ocular reflex neurons, connectivity and subtype are inextricable: eyes-down motor neurons must synapse onto eyes-down muscles, and nose-up afferents must receive input from up-tuned hair cells. Thus, synapse formation – even originating from stochastic connectivity – could define subtype^118^. Importantly, however, birthdate differentiates the subtype of vestibular projection neurons^119^, extraocular motor neurons^8^, and sensory afferents^55^ all *before* they connect to each other. Further, projection neurons and extraocular motor neuron soma differentiate coincidentally and project early, and synaptogenesis is delayed until somatic differentiation is complete across both nuclei (Figure 6). This argues that fate and assembly in the vestibulo-ocular reflex circuit follows from time, not connectivity.

Is time *deterministic* for vestibulo-ocular reflex circuit assembly? Across the entire vestibulo-ocular reflex circuit, the nose-up/eyes-down channel develops earlier than its nose-down/eyes-up counterpart (Figure 7). The different embryological origins of channel components (ear, brain, and eye) suggest that a broad signal is required to coordinate circuit assembly. Such global timing mechanisms are well-established in *Drosophila* nervous system^120,121^. Recent profiling studies have identified temporally-specific, though spatially-broad, patterns of gene expression in vertebrates^122^. Temporal and/or spatial transplantation approaches^4,123^ offer a means to test this hypothesis. We propose that the zebrafish vestibulo-ocular reflex circuit is a powerful model to test if global timing mechanisms coordinate channel-specific synaptic assembly to enable a functional sensorimotor circuit.

### Temporal influences on sensorimotor circuit development

We propose vestibulo-ocular reflex circuit development more closely resembles invertebrate circuit development than vertebrate sensorimotor circuits. For invertebrates, the molecular logic that links time, fate, and circuit-level connectivity are well-established^34–37^. There, time reflects the synchronized and sequential expression of specific transcription factor codes, pre-mitotically and independently of space^86,124^. In vertebrates, such tight links between lineage and fate are undefined. Though relationships between time and circuit function have been demonstrated in vertebrate circuits –for example, in zebrafish locomotor circuits for swimming^45,82,125^ –even there, fate and connectivity are thought to arise from post-mitotic spatial cues^97,126,127^. In our system, the rough topography but clear temporal delineation among projection neuron subtypes argues against the primacy of spatial cues in vestibulo-ocular reflex circuit assembly.

We discovered unexpected spatiotemporal organization among vestibular projection neurons, their sensory inputs, and their motor outputs. The spatial topography we observe challenges previous reports of “disorganization” among brainstem vestibular neurons, instead illustrating that balance nuclei can resemble other sensory areas. As topographic maps reflect developmental pressures, our findings illuminate the mechanisms that may govern projection neuron fate and reflex circuit assembly, a key step forward in understanding how balance circuits come to be. Finally, the contemporaneous development of vestibulo-ocular reflex channels links vertebrate circuit assembly to temporal principles established in invertebrates. Though later maturation may disrupt the topography we observe in the larval zebrafish, our discoveries and inferences nevertheless illustrate how time may shape functional vertebrate sensorimotor circuits.

## ACKNOWLEDGMENTS

Research was supported by the National Institute on Deafness and Communication Disorders of the National Institutes of Health under award numbers R00DC012775, R56DC016316, R01DC017489, and F31DC019554, and the National Institute of Neurological Disorders and Stroke under award numbers F99NS129179, T32NS086750, UF1NS108213, and U01NS094296. The authors would like to thank Başak Sevinç and Hannah Gelnaw for assistance with fish care, and Jeremy Dasen, Claude Desplan, Robert Froemke, Michael Long, Katherine Nagel, Dan Sanes, along with the members of the Schoppik and Nagel labs for their valuable feedback and discussions.

## AUTHOR CONTRIBUTIONS

Conceptualization: DG and DS, Methodology: DG, KH,VV, CP, KP, WL, EH, MB, MG, and DS, Investigation: DG, SH, MB, Visualization: DG, Writing: DG, Editing: DS, Funding Acquisition: DG and DS, Supervision: DS.

## STAR METHODS

### RESOURCE AVAILABILITY

#### Lead Contact

Further information and requests for resources and reagents should be directed to and will be fulfilled by the lead contact, David Schoppik.

#### Materials Availability

This study did not generate new unique reagents.

#### Data and code availability

All data and code used in this paper have been deposited at the Open Science Framework, DOI 10.17605/OSF.IO/MRG5C

### EXPERIMENTAL MODEL AND SUBJECT DETAILS

#### Fish care

All protocols and procedures involving zebrafish were approved by the New York University Langone School of Medicine Institutional Animal Care & Use Committee (IACUC). All larvae were raised at 28.5°C at a density of 20-50 larvae in 25-40 ml of buffered E3 (1mM HEPES added). Larvae used for photofill and birthdating experiments were raised in constant darkness; all other fish were raised on a standard 14/10h light/dark cycle. Larvae for experiments were between 1-5 days post-fertilization (dpf).

#### Transgenic lines

Experiments were conducted on the mifta -/- background to remove pigment. Photofill experiments were conducted on the *Tg(-6.7Tru.Hcrtr2:GAL4-VP16; UAS-E1b:Kaede)* background^28,57,58^. Photofill localization at nIII/nIV was validated with the *Tg(isl1:GFP)* line^128^. Two-photon calcium imaging experiments were performed on the *Tg(-6.7Tru.Hcrtr2:GAL4-VP16; UAS:GCaMP6s; atoh7:gap43-RFP)* background^129^. Functional birthdating experiments used larvae from the *Tg(-6.7Tru.Hcrtr2:GAL4-VP16; UAS:Kaede; UAS:GCaMP6s; atoh7:gap43-RFP)* background. SCAPE experiments used blind larvae from the *Tg(-6.7Tru.Hcrtr2:GAL4-VP16; UAS:Kaede; UAS:GCaMP6s; atoh7:gap43-RFP; lakritz^th241^)* background^130^. All larvae used were selected for brightness of fluorescence relative to siblings. Mendelian ratios were observed, supporting that selected larvae were homozygous for a given allele. To assay loss of the utricular otolith, we used the rks^vo66/vo66^ *rock solo* otogelin knockout^68^. To label the VIII^th^ nerve, we used *Tg(-17.6isl2b:GFP)*^131^.

### METHOD DETAILS

#### Confocal imaging

Larvae were anesthetized in 0.2 mg/mL ethyl-3-aminobenzoic acid ethyl ester (MESAB, Sigma-Aldrich E10521, St. Louis, MO) prior to imaging except where noted. Larvae were mounted dorsal side-up (axial view) or lateral side-up (sagittal view) in 2% low-melting point agarose (Thermo Fisher Scientific 16520) in E3. Images were collected on a Zeiss LSM800 confocal microscope with a 20x water-immersion objective (Zeiss W Plan-Apochromat 20x/1.0). All imaging windows were 266×133 μm. Anatomy stacks of the tangential vestibular nucleus spanned approximately 50 μm in depth, sampled every micron. Raw image stacks were analyzed using Fiji/ImageJ^132^.

#### Retrograde photolabeling of ascending projection neuron somata

Retrograde photolabeling experiments were performed to define the anatomical boundaries of projection neuron somata and generate a reference spatial framework (Methods, “Spatial registration of imaged neurons”). Ascending projection neurons from the tangential vestibular nucleus were visualized using the *Tg(-6.7Tru.Hcrtr2:GAL4-VP16; UAS-E1b:Kaede)* line. Kaede is a photolabile protein that irreversibly converts from green to red with ultraviolet light^56^. To prevent background photoconversions, all larvae were raised in darkness. Photoconversions were performed using a confocal microscope (Zeiss LSM800) 5 dpf larvae were anesthetized and mounted axially in 2% low-melt agarose. Ascending projections were identified using the ascending bundle of axonal arbors off the medial longitudinal fasciculus at nIII/nIV^28^. Localization was validated in separate experiments on the Tg(isl1:GFP) background, which labels nIII and nIV ocular motor neurons^128^. An imaging window was centered on a single hemisphere over the midbrain-hindbrain boundary between nIII and nIV, just lateral to the medial longitudinal fasciculus. The area was repeatedly scanned with a 405 nm laser for 20-30 seconds until fully converted. Immediately after, the same plane was imaged to assess for off-target photoconversion (e.g., conversion of axonal bundles lateral to the medial longitudinal fasciculus). Larvae were unmounted and left to recover in E3 in darkness for approximately four hours to permit converted (red) Kaede to retrogradely diffuse to the somata. Photofilled fish were then re-anesthetized and mounted laterally and then axially. An imaging window was centered over hindbrain rhombomeres 4-6. Photofilled somata were identifiable by their center-surround fluorescence appearance: red converted cytoplasm, surrounding a green unconverted nucleus. Retrograde photolabeling experiments were performed in N = 3 larvae.

#### Optical tagging of neurons by birthdate

Neuronal somata and axons were optically tagged by their time of terminal differentiation using whole-embryo Kaede photoconversions^56^. All larvae were raised in darkness to prevent background photoconversion. Whole embryos were photoconverted for five minutes at experimenter-defined timepoints. Photoconversions were performed using a custom-built apparatus with a 405 nm LED bulb. At 5 dpf, anesthetized larvae were imaged on a confocal microscope. Only neurons born before the time of photoconversion contained converted (red) Kaede. These neurons sometimes also contained unconverted (green) Kaede, reflecting the continued production of new Kaede protein between the time of conversion and the time of imaging. Neurons that were born after the time of conversion contained only green (unconverted) Kaede. For each experiment, basal conversion was estimated by imaging control larvae raised in darkness until confocal imaging. Conversion was not observed in these larvae.

#### Somatic birthdating experiments

Whole embryos were birthdated at an experimenter-defined developmental stage: 22, 24, 28, 30, 33, 36, 42, or 48 hpf. Each embryo was converted at only one timepoint. n=5 hemispheres from at least N=3 separate larvae were analyzed for each timepoint between 22-42 hpf, and n = 3 hemispheres from N=3 larvae were analyzed for the 48 hpf timepoint. For each timepoint, we quantified the ratio of the number of converted neurons to the total number of neurons in that hemisphere.

#### Tonic and impulse pitch-tilt stimuli

All experiments were performed on 5 dpf larvae (Transgenic lines). Stimulation paradigms were presented as previously reported in^54^ and are described again below. Larvae were paralyzed using bath application of 0.8 mg/mL pancuronium bromide (Sigma-Aldrich P1918, St. Louis, MO) for 8-10 minutes. Larvae were then mounted dorsal-up in 2% low-melt agarose in E3 in the center of the mirror on a large beam diameter single-axis scanning galvanometer system (ThorLabs GVS011, 10 mm 1D). The driver board was set to a mechanical position signal input factor of 0.5 V per degree, for a maximum mechanical scan angle of ±20.0°. Inputs of ±9.5 were delivered to the galvanometer to drive ±19°rotations in the pitch (up/down) axis.

Tonic pitch-tilt stimuli were delivered over a 65-second period. The stimulus began with an initial five-second baseline at a horizontal (0°) imaging plane, with the tangential vestibular nucleus in view. Subsequently, an instantaneous (4 ms) step was delivered to rotate the larvae −19°away from horizontal. This eccentric position was held for 15 seconds before delivering a (4 ms) horizontal restoration step, held again for 15 seconds. This process was then repeated for the nose-up position (+19°). Nose-down (−19°) stimuli were always presented first, followed by the nose-up (+19°) rotation. A neuron’s directional response was defined as the change in GCaMP6s fluorescence in the first second of the return to horizontal. To control for presentation order effects, separate experiments with an inverted presentation order were performed on one larva (n=66 neurons). No significant differences were observed between these responses and all other neuronal responses.

Impulse rotations were delivered over a similar 65-second period. Larvae were presented with two sets of impulse stimuli. Each impulse was 10 ms in duration. The first impulse set contained a 4 ms downwards (−19°) rotation, a 2 ms hold,and a 4 ms restoration step to horizontal. The second impulse step contained a similar 4 ms rotation upwards (+19°), a 2 ms hold, and then a 4 ms restoration step. Each impulse was followed by a 30-second imaging window at horizontal. Impulse delivery was presented just before the equivalent restoration to horizontal in the tonic stimulus (20 and 50 seconds, respectively) to facilitate comparison of the impulse contribution to the tonic restoration step.

Tonic and impulse rotations were presented in alternating stimulus sets. Impulse rotations were always presented first, followed by tonic rotations. Three stimulus repeats (six total trials) were presented for each imaged plane for two-photon experiments, and ten trial sets (20 total trials) for SCAPE experiments.

#### Two-photon calcium imaging

Two-photon calcium imaging experiments were conducted using a 20x water immersion objective (Olympus XLUMPLFLN20xW 20x/1.0). The GCaMP6s calcium indicator was excitepd using an infrared laser (SpectraPhysics MaiTai HP) at 920 nm using approximately 6.1-18.8 mW of power, measured at the specimen with a power meter (ThorLabs S130C). Stimulus imaging was conducted using ThorLabs LS 3.0 software. All experiments took place in the dark

Sighted 5 dpf larvae were paralyzed and mounted dorsal-up on a large beam diameter galvanometer as described above. Imaging windows were centered over rhombomeres 4-6 with both the tangential vestibular nucleus and the Mauthner neuron in view. In larvae on the *Tg(isl1:GFP)* background, GFP brightness oversaturated the photomultiplier tube in ventral planes. For these experiments, each hemisphere was imaged separately, and ventral planes were imaged with higher magnification (112×68 μm imaging window) than dorsal planes (148×91 μm imaging window). All other larvae were imaged using 205×127 μm imaging windows, with both hemispheres of the tangential vestibular nucleus in view.

An anatomical image stack of the tangential nucleus was acquired for each fish at 1 frame/second (5.2 μs pixel dwell time). Anatomy stacks spanned approximately 50 μm in depth, sampled every micron. These stacks were used to determine stimulus imaging planes. Efforts were made to sample the entire nucleus for each fish. Typically, this required sampling 6-10 planes, spaced 3-6 μm apart based on cell density. The brightness of the calcium indicator increased from the ventral-to dorsal-most neurons in a highly-stereotyped manner. Laser power was correspondingly adjusted for each plane, such that greater power (18.8 mW) was used for ventral planes compared to dorsal (6.1 mW) planes. To control for potential photobleaching effects, the nucleus was always sampled from the ventral to dorsal direction. All stimulus imaging experiments were performed at 3 frames/second (2.2 μs pixel dwell time). Total experiment time lasted approximately two hours per fish.

#### Otogelin mutant imaging and analysis

Experiments were performed on sighted 5 dpf larvae (Transgenic lines). Null otogelin mutants and sibling controls (non-phenotypic) were identified by the absence or presence of the utricular otoliths, respectively, at 5 dpf. Stimuli were delivered to three null mutants and three sibling controls under a two-photon microscope as previously described. Projection neuron responses were analyzed as previously described. 11 neurons (mutant: n=3 neurons; control: n=8 neurons) were excluded due to technical issues in imaging, leaving 183 neurons from null mutants and 144 neurons from sibling controls for further analysis (n=63±9 neurons per mutant fish, n=54±2 neurons per sibling fish, across both hemispheres). All neurons were analyzed, regardless of whether they exhibited a significant response to either stimulus. Responses were quantified as the distribution of the maximum change in fluorescence to the preferred (tonic) direction. Significant differences between the null and control distributions were assessed using a two-sample, one-tailed KS test.

#### VIII^th^ nerve lesions, imaging, and analysis

Experiments were performed on a two-photon microscope using sighted 5 dpf larvae (Transgenic lines). Projection neurons were imaged before and after uni-lateral lesions of both the anterior and posterior semicircular canal branches of the VIII^th^ nerve. To obtain a pre-lesion baseline, paralyzed larvae were mounted dorsal-up (axially) and presented with tonic and impulse stimuli as described previously. Larvae were then re-mounted lateral side-up (sagittally) to visualize the VIII^th^ nerve. Lesions were targeted to the anterior and posterior semicircular canal branches of the VIII^th^ nerve, approximately 20-30 μm from the cristae. Lesions were performed using a pulsed infrared laser (SpectraPhysics Spirit 8W) at 1040 nm (400 fs pulse duration, 4 pulses per cell over 10 ms) at 25-75 nJ per pulse. Larvae were immediately imaged to confirm the extent of the lesion. Larvae were left to recover for ten minutes, re-mounted axially, and then imaged again following tonic and impulse stimuli presentation.

Neurons were only analyzed if they were identifiable in both pre- and post-lesion imaging. This left us with 62 control neurons (N=4 larvae) and 56 lesioned neurons (N=4 larvae). For impulse response analyses, neurons were only analyzed if they were impulse-responsive in the pre-lesion imaging condition (n=29 control neurons, n=33 lesioned neurons). Analyses were performed on the maximum change in fluorescence in the first second of the response following tonic and impulse rotations. All statistical analyses were conducted using a one-tailed, two-sample KS test.

#### Functional birthdating

Experiments were performed on sighted larvae (Transgenic lines). Embryos were birthdated at 30, 36, 42, or 48 hpf as previously described. At 4 dpf, larvae were imaged on a confocal microscope to identify neurons that were born by the time of photoconversion (red, converted Kaede; early-born) or born after the time of photoconversion (green, unconverted Kaede only; late-born). Notably, late-born (green Kaede) neurons were clearly distinguishable from GCaMP6s-positive neurons given the differential localization of each fluorophore (whole-cell vs. nucleus-excluded, respectively). To avoid possible effects of an anesthetic on subsequent measures of calcium activity, larvae were not anesthetized for confocal imaging. We validated that the spatial distributions of projection neurons were constant between 4 dpf and 5 dpf by mapping neurons from five non-birthdated reference stacks from each age. No significant differences were observed between these distributions. Larvae were left to recover overnight in E3 in normal light/dark conditions.

Two-photon calcium imaging was performed at 5 dpf on paralyzed larvae. To spectrally separate green GCaMP6s from unconverted (green) Kaede, larvae were photoconverted for five minutes on our 405 nm LED apparatus to ubiquitously convert all Kaede to red. Tonic and impulse rotations were presented as previously described. Two channels (green and red) were imaged to facilitate localization of Kaede-positive neurons. Only Kaede-positive neurons were analyzed further. We manually registered the confocal stack for each imaged hemisphere to our reference framework as previously described. The coordinates and birth status (born before/born after the time of conversion) were recorded for each neuron. Neurons were then identified in the appropriate sampled stimulus plane. This permitted alignment of a given neuron’s spatiotemporal developmental properties with its functional identity and rotation response features. We analyzed a total of n = 74 neurons from N=10 larvae birthdated at 30 hpf; n = 376 neurons from N = 7 larvae birthdated at 36 hpf; n=264 neurons from N=5 larvae birthdated at 42 hpf; and n=41 neurons from N=10 larvae birthdated at 48 hpf.

#### Swept, confocally-aligned planar excitation (SCAPE) calcium imaging and analysis

Volumetric calcium imaging experiments were performed using a custom-built SCAPE 2.0 microscope design ^49^ with a 20x water immersion objective (Olympus XLUMPLFLN20xW 20x/1.0). Experiments were performed on blind 5 dpf larvae (Transgenic Lines). Paralyzed larvae were mounted dorsal-up (axial) on a large beam diameter galvanometer and imaged in an oblique (approximately 26°) coronal view. A 179×896 μm imaging window was centered over the tangential vestibular nucleus. Each volume spanned approximately 50 μm in the rostrocaudal axis, sampled every 3 μm. The rostral- and caudal-most extent was identified using the facial nucleus (nVII) and the Mauthner neuron, respectively. A high-resolution anatomical image stack was acquired at a rate of approximately 1 frame/second (0.0066 volumes per second), while stimulus imaging was performed at a rate of approximately 130 frames/second (5 volumes per second).

Larvae were presented with ten stimulus repeats (20 total trials) of the tonic and impulse stimuli as previously described. Two modifications were made to this stimulus: i) the initial baseline imaging period was extended to 15 seconds, and ii) nose-up and nose-down tilt presentation order was randomized each trial. The total experiment time per fish was approximately one hour. A second high-resolution anatomical image stack was acquired after stimulus imaging to assess for photobleaching effects. No significant photobleaching was observed.

#### Electron microscopy

Serial section electron microscopy images were obtained as described ^55^. Briefly, an ultrathin (60 nm) section library of the entire zebrafish head was initially imaged at 18.8×18.8×60 nm^3^ per voxel and 56.4×56.4×60 nm^3^ per voxel. All myelinated axons, including those from utricular and semicircular canal afferents, were reconstructed ^133^. A volume on the right side of the head, including the utricular hair cells, afferents, and most of the vestibular brainstem, was later reimaged at 4.0 Œ 4.0 Œ 60 nm^3^ per voxel, allowing identification of synaptic contacts between vestibular afferents and their brainstem targets.

Projection neurons of the tangential nucleus were identified by three criteria: soma position caudal to the vestibular nerve; axonal projections that cross the midline and begin to ascend ^27^; and the presence of synaptic contacts from either utricular or canal afferents. Several putative projection neurons are not included here because their axons, which exit the high-resolution reimaged territory, could not be followed adequately; these neurons are described elsewhere^134^. Therefore, the dataset is biased towards early-myelinated tangential neurons.

Each projection neuron, utricular afferent, and canal afferent was reconstructed as fully as possible. Canal afferents exited the high-resolution reimaged volume, and therefore could only be fully reconstructed if they were myelinated, whereas all utricular afferents were reconstructed. Onto projection neurons, there were no contacts from horizontal (medial) canal afferents, 89 synaptic contacts from anterior or posterior canal afferents, and 104 contacts from utricular afferents. Directional tuning of canal afferents was assumed based on canal identity (anterior canal: nose down; posterior canal: nose up). Directional tuning of utricular afferents was computed as a weighted vector sum of the hair cells forming ribbon synapses with each afferent^55^.

#### Axon birthdating

Birthdating was performed as previously described. Whole embryos were converted at 36 hpf. At 3 dpf, anesthetized larvae were mounted lateral side-up (sagitally) to facilitate visualization of the dorsoventral separation between axon tracts. Axons were identified as dorsal-projecting based on location in the medial longitudinal fasciculus, and if they exhibited the characteristic bundle of axonal arbors off of nIV^28^. We analyzed at least one hemisphere from three separate larvae.

### QUANTIFICATION AND STATISTICAL ANALYSIS

#### Classification and analysis of two-photon calcium responses to tonic and impulse rotations

We sampled 153 planes from 22 larvae. Sampling planes were pre-processed using Fiji/ImageJ ^132^. For each plane, polygonal regions of interest (ROI) were drawn around all projection neuron somata visible in a maximum intensity projection of the first stimulus trial. To correct for minor (1-2 μm) inter-trial movement, ROI positions were manually adjusted in the XY axes. 37 planes were excluded due to excessive movement that caused more than one neuron to appear within an ROI. Raw fluorescence traces from polygon ROIs were extracted using Matlab R2020b (MathWorks, Natick, MA, USA) and normalized by ROI size to account for variation in imaging window size. A neurons response was defined as the fluorescence values over the 15-second horizontal imaging window following an eccentric rotation.

To standardize responses across stimulus repeats, neurons, and larvae, all fluorescence responses were normalized by a baseline period. Some fluorescence responses remained elevated above baseline at the end of an imaging window; we accounted for this elevation in the following way, with post-hoc validation: The baseline for the static nose-down response was defined as the mean fluorescence value of the initial five-second horizontal imaging window for that trial. The baseline for the static nose-up response was defined as the mean fluorescence value of the last three seconds of the nose-down response. For impulse rotations, the baseline was defined as the mean fluorescence value of the five seconds immediately preceding the rotation. Change in fluorescence (ΔF/F) was quantified by subtracting the appropriate baseline from each frame of the response and then normalizing by the appropriate baseline. To validate that using different baseline periods did not account for differential nose-up and nose-down response properties, all analyses were repeated for the n=66 neurons presented with an inverted directional rotation order. These analyses did not significantly change any conclusions.

We analyzed the calcium responses of 694 unique neurons. 120 neurons were excluded due to technical issues encountered during imaging. This included larvae becoming loose from the agarose, causing excessive, non-stimulus locked positional movement and response jitter. This also included loss of water between the specimen and the objective, which caused significant decreases in fluorescence brightness between trials due to the air/water interface. We excluded an additional 57 dorso-caudal cells located outside the spatial bounds of the nucleus, as defined from retrograde photofill experiments. This left us with 517 neurons for further analysis. Each neuron’s raw fluorescence and ΔF/F timecourses of the tonic and impulse responses were manually inspected. Neurons exhibited one of three response patterns to tonic tilts: i) transient excitation that peaked in the first second and then decayed, ii) recovery from suppression, which frequently increased to an above-baseline value, or iii) no response. Only transient excitation was observed in response to impulse stimuli. We manually classified each neuron’s response pattern to each directional stimulus. To control for bias in manual classification, we only analyzed excitatory transients further if, for at least two trials, the mean raw fluorescence response in the first second was at least two standard deviations greater than baseline. 36 neurons produced clear excitatory transient responses with peaks below baseline, likely due to high background noise, and were excluded from further analysis. All subsequent analyses were conducted on the mean ΔF/F response across all stimulus repeats.

We assigned a nose-up or nose-down identity based on the direction that produced stronger response. To quantify the strength of directional tuning, we defined a directionality index. The directionality index normalized the difference in ΔF/F responses to the up and down rotations by their sum. Thus, negative directionality index values represented stronger nose-down responses (maximum of - 1), positive values indicated stronger nose-up responses (minumum of - 1), and a value of zero indicated equal nose-up and nose-down responses. To distinguish neurons with a clear directional preference from those with no preference, we set a qualitative threshold at 0.1, which represents a 22% or greater difference in relative response strength. All conclusions remained constant when eliminating or doubling this threshold. Here, 10 neurons were excluded because they produced inconsistent directional responses across stimulus repeats.

We quantified several additional response properties. The coefficient of variation across stimulus repeats was quantified as a ratio of the mean ΔF/F response across stimulus repeats to their standard deviation. Response strength was quantified using two metrics: the mean and integral of the ΔF/F response in the first second after the return to horizontal. The integral was estimated using the trapezoidal method with unit spacing. Response mean and integral were highly correlated (r=0.9971; p<0.001); therefore, only response mean was used for further analyses. Neuron response strength was illustrated by normalizing each neuron’s mean ΔF/F response against the maximum ΔF/F response observed for that subtype. Lastly, to estimate the proportion of the tonic response that may originate from the rapid (4 ms) restoration step, we subtracted the impulse ΔF/F response from the tonic ΔF/F response to generate a “residual.” To calculate the fraction of the tonic response that may be explained by the impulse step, we quantified the ratio of the residual response to the original tonic response. Lastly, some neurons exhibited slight directional tuning to impulse rotations (mean directional strength = 0.30±0.26). Therefore, impulse responses to each impulse set (up/down, then down/up) were analyzed separately.

#### Spatial registration of imaged neurons

All imaged neurons were manually registered to a reference coordinate framework in Adobe Illustrator (2021). The reference framework was generated from the maximum intensity projections (MIPs) of three retrogradely-photofilled tangential vestibular nuclei, imaged on a confocal microscope. MIPs were aligned using stereotyped anatomical landmarks such as the Mauthner neuron cell body, the medial longitudinal fasciculus, and the otic capsule. MIPs for all anatomical image stacks were manually registered to this reference framework. To control for potential bias in manual registration, all registrations were verified by two independent observers (D.G. and S.H.). To control for differences in digital magnification across microscopes and larvae, all stacks were uniformly scaled to the resolution of the reference stacks (4.8 pixels/ μm).

Empirically, we observed that projection neuron somata were ~5 μm in diameter. To simplify localization, each anatomical image stack was uniformly subdivided into eight, 5 μm thick (dorsoventral) planes, such that each neuron would appear in no more than two planes. Subdivisions were assessed for consistency using distinct anatomical landmarks. Ventral planes (z3-z5) always included the Mauthner cell body. The dorsoventral midpoint (z4-z5) appeared in the same plane as the mediolateral midpoint: dorsal neurons were exclusively medial to the midpoint, and ventral neurons were exclusively lateral. The dorsomedial-most neurons in the core of the nucleus were always present in the second-most dorsal plane (z7). All subdivisions were verified by two independent observers (D.G. and S.H.). Each dorsoventral subdivision was manually registered to the MIP for that nucleus.

Neurons were manually localized to a dorsoventral subdivision based on the plane in which the soma fluorescence was brightest (soma center). Reference neurons, represented as circles approximately the diameter of a neuron, were centered over the soma in the appropriate subdivision in Illustrator. The dorsoventral subdivision and the rostrocaudal (X) and mediolateral (Y) Illustrator coordinates were manually recorded for each neuron. For standardization, their XY coordinates were normalized by subtraction from an absolute (0,0) point. This point was defined as the upper left corner of a rectangular bound, which was drawn over the extent of the photofilled tangential vestibular nucleus. Standardized coordinates were then used to recreate a spatial map of neurons imaged across all fish in Matlab (R2020b), and to link a given neuron’s spatial coordinates with its functional properties. 363/467 imaged neurons were mapped.

#### Statistical analysis of spatial organization

Spatial organization was evaluated in two ways: i) across the entire tangential vestibular nucleus, and ii) with respect to each spatial axis separately. Global organization (three-dimensional coordinates) was evaluated using a one-way, multivariate analysis of variance test. Organization in each individual axis was evaluated using a two-tailed, two-sample Kolmogorov-Smirnov (KS) test. To assess whether spatial organization was due to chance, we randomly permuted the feature of interest (e.g., up/down subtype) for all neurons while preserving the observed occurence of that feature and spatial coordinates. 100 permuted distributions were generated and statistical analyses repeated. No significant effect was ever observed in permuted distributions.

Qualitatively, we observed that the spatial separation between nose-up/nose-down subtypes was clearest in a pseudo-sagittal view of the tangential vestibular nucleus, though all confocal and two-photon images were acquired in an axial view. To determine the optimal sagittal view, we tested a set of affine transformations. 26,571 combinations of azimuth and elevation values were tested. The optimal combination (azimuth = −26.7, elevation = 51.9) was defined as that which maximized the spatial separation between nose-up and nose-down subtypes across all three spatial axes, per two-sample KS tests. All reconstructions of birthdated and two-photon neuron soma position are shown as affine-transformed coordinates. All statistical analyses were performed on raw (non-transformed) coordinates.

#### Analysis of SCAPE images

SCAPE images were analyzed on a per-pixel basis. For each pixel, we computed the mean change in fluorescence of the first second (5 volumes) of each stimulus response relative to the initial 15-second baseline period for a given trial. The mean change in fluorescence was averaged across all ten stimulus repeats for the tonic and impulse stimuli separately. A directionality index was computed as previously described. Each pixel was then pseudo-colored according to the direction that evoked the greater change in fluorescence, as determined by the directionality index. Color intensity was scaled such that the maximum intensity corresponded to the maximum directionality index value for that subtype.

To mirror the sagittal orientation of the two-photon and confocal datasets, SCAPE images were rotated 90°around the Y-axis with interpolation. To correct the oblique angle of the SCAPE laser, an affine transformation was performed using the TransformJ plugin^135^ in ImageJ and the following transformation matrix:

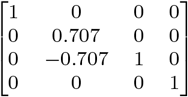

#### Quantification of post-synaptic axonal varicosity growth

Axo-somatic varicosities were quantified as a proxy for synapses^73^. Motor neuron soma were classified as dorsal or ventral based on their position relative to the appearance of nIV^8^. Anesthetized larvae were mounted dorsal-up and imaged on a confocal microscope between 50-90 hpf. Larvae were imaged no more than three times to minimize photobleaching. Varicosities were qualitatively defined as 1 μm axo-somatic spheres. We observed approximately three times as many varicosities onto all dorsal motor neurons than to ventral neurons, in line with the differential densities of nIII and nIV^8^. Therefore, for each timepoint, varicosity growth was quantified as a percentage of the mean number of varicosities observed at 90 hpf.

---

**Figure S1:**
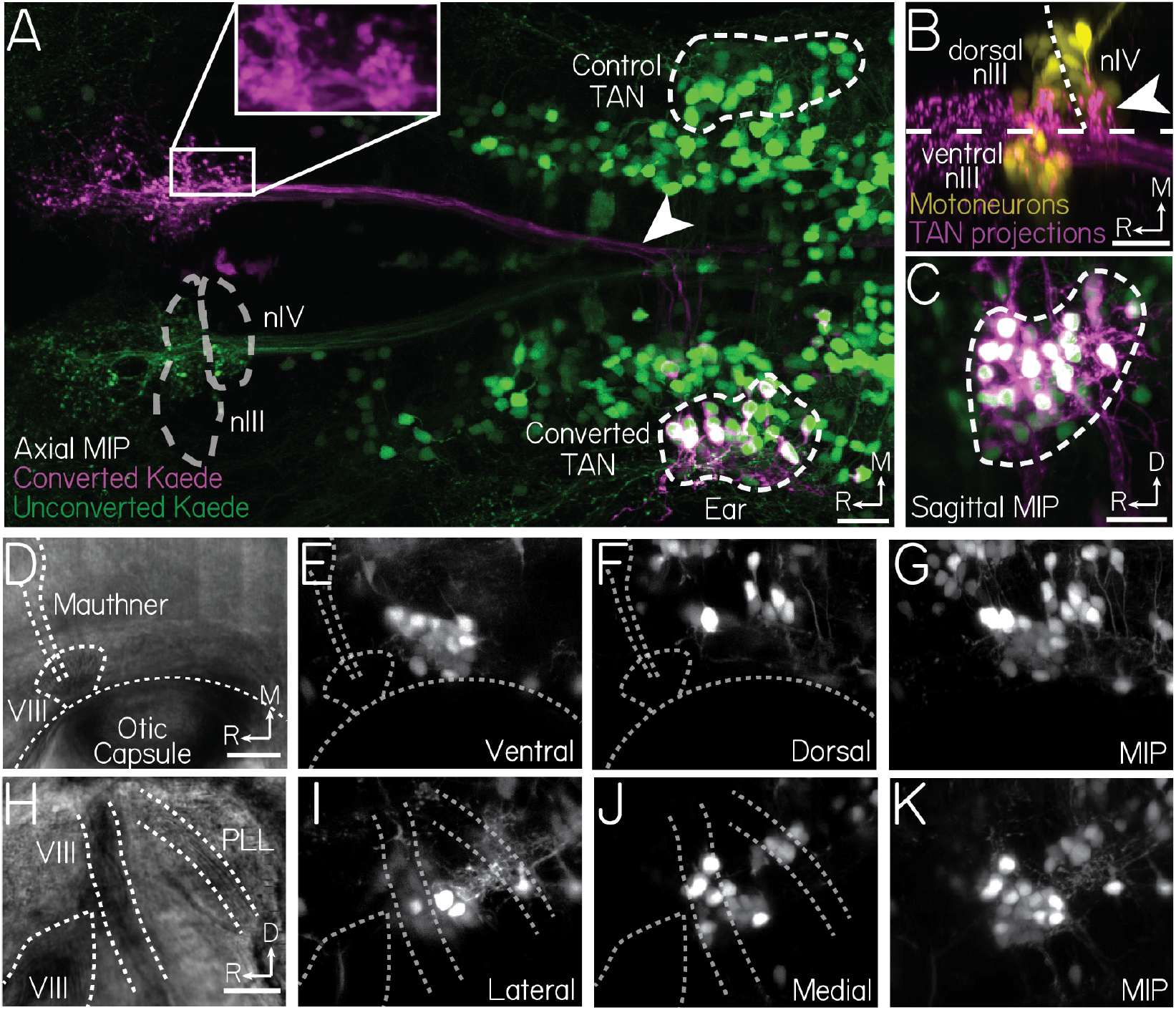
Localization of the tangential vestibular nucleus, Related to Figure 1. (A) Retrograde photofill technique. Maximum intensity projection (MIP) of the tangential vestibular nucleus and its ascending projections at 5 dpf. White box shows conversion target region. White arrow points to ascending projections. Grey dashed lines outline the position of the extraocular motor nuclei. White dashed lines outline the bounds of photofilled soma in the tangential vestibular nucleus. (B) Photoconversion target region. White arrow points to the characteristic axonal arborizations at cranial nucleus IV (nIV). Motor neurons labeled in yellow. Ascending projections labeled in magenta. Horizontal dashed line indicates midline. Diagonal white dashed line shows midbrain-hindbrain boundary. (C) Center-surround appearance of converted (magenta) and unconverted (green) Kaede in retrogradely-labeled soma. (D-G) Location of the tangential vestibular nucleus in an axial view. (D) Transmitted light view of the tangential vestibular nucleus, 15 μm ventral to the base. Grey dashed lines outline the otic capsule, VIII^th^ cranial nerve, and lateral dendrite of the Mauthner neuron. (E) Ventral plane, approximately 15 μm dorsal to the base. Landmarks in D labeled as grey dashed lines. (F) Dorsal plane, approximately 35 μm dorsal to the base. (G) Maximum intensity projection. (H-K) Location of the tangential vestibular nucleus in a sagittal view. (H) Transmitted light view, 10 μm from the lateral edge. White dashed lines outline the entry point and ascending branch of the VIII^th^ nerve (VIII) and the posterior lateral line branch (PLL). (I) Lateral plane, approximately 10 μm medial to the lateral edge. (J) Medial plane, approximately 30 μm medial to the lateral edge. (K) Maximum intensity projection. All scale bars 20 μm.

**Figure S2:**
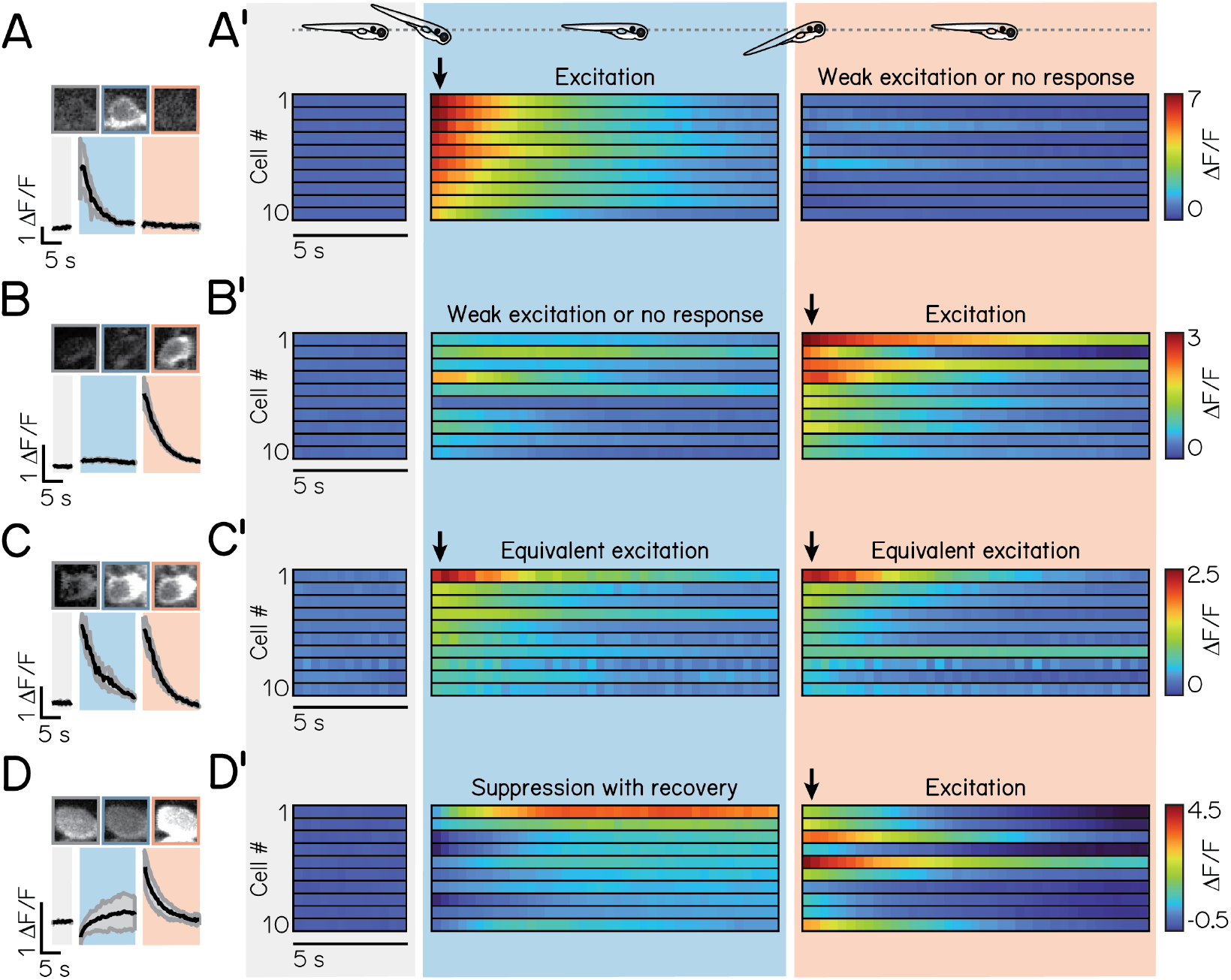
Response profiles of tangential vestibular neurons to tonic pitch-tilt rotations, related to Figure 2. (A) Example response from a nose-down neuron with no activity to the non-preferred direction. Black line indicates mean response across three stimulus repeats; shaded bars show standard deviation. Same example neuron as in Figure 2D. (A’) Heatmap showing the timecourse of calcium activity for n=10 example nose-down neurons with no response or weak excitation to the non-preferred (nose-up) direction. Each row represents a distinct neuron. Shaded bars indicate time at horizontal for baseline (grey), following the nose-down rotation (blue), and following the nose-up rotation (orange). Black vertical arrow points to excitatory responses used to assign a nose-up, nose-down, or no selectivity identity. (B) Example response from a nose-up neuron with no activity to the non-preferred direction. Same example neuron as in Figure 2D. (B’) Heatmap showing the timecourse of calcium activity for n=10 example nose-up neurons with no response or weak excitation to the non-preferred (nose-down) direction. (C) Example response from neuron with no directional selectivity. Same example neuron as in Figure 2D. (C’) Heatmap showing the timecourse of calcium activity for n=10 example neurons with no directional selectivity. (D) Example response from nose-up neuron that exhibits suppression with recovery to the non-preferred (nose-down) direction. (D’) Heatmap showing the timecourse of calcium activity for n=10 example nose-up neurons that exhibit suppression with recovery to the non-preferred direction.

**Figure S3:**
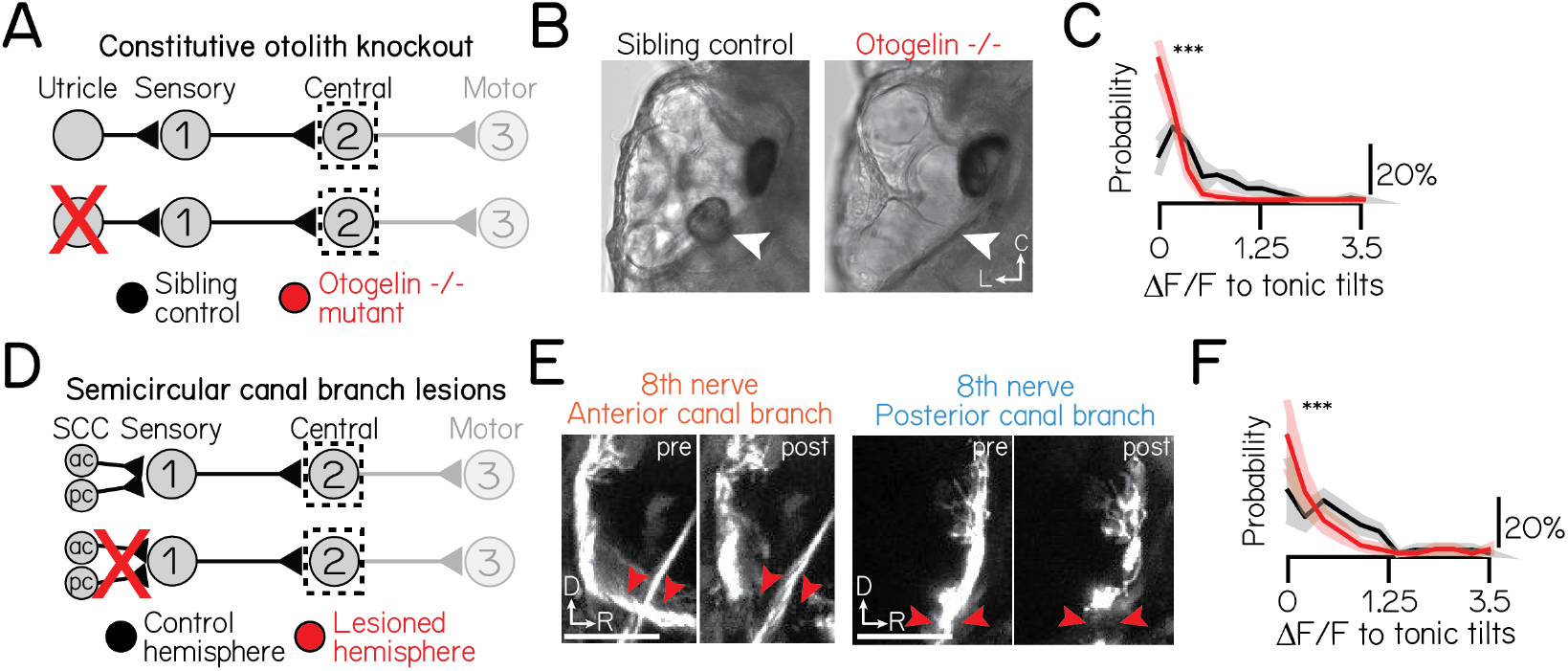
Responses to tonic pitch-tilt rotations originate primarily from the utricular otoliths, Related to Figure 2. (A) Circuit schematic for utricular otolith knockout experiments. Black dashed lines outline the tangential nucleus as the circuit element of focus. (B) Example images from a control larvae (left) and an otogelin null mutant (right). White arrow points to the position of the utricular otolith. (C) Probability distributions of the maximum ΔF/F response to preferred directional pitch-tilts in control (black) and otogelin mutants (red). Solid lines shows mean from jackknife resampling; shaded bars, standard deviation. Controls: n=144 neurons, N=3 fish; Mutants: n=183 neurons, N=3 fish. (D) Circuit schematic for uni-lateral VIII^th^ nerve lesions (anterior and posterior semicircular canal branches). (E) Example images from larvae before and after uni-lateral VIII^th^ nerve lesions. Left and right image sets show the anterior and posterior semicircular canal branches, respectively, before and after lesion. Both branches are lesioned in each experiment. Red arrows point to lesion sites. (F) Probability distributions of the maximum ΔF/F response to preferred directional pitch-tilts in control (black) and lesioned (red) hemispheres. Solid lines shows mean from jackknife resampling; shaded bars, standard deviation. Control hemisphere: n=57 neurons, N=4 fish; Lesioned hemisphere: n=49 neurons, N=4 fish. All scale bars, 20 μm. All data: Three stars denotes significance at the p<0.001 level.

**Figure S4:**
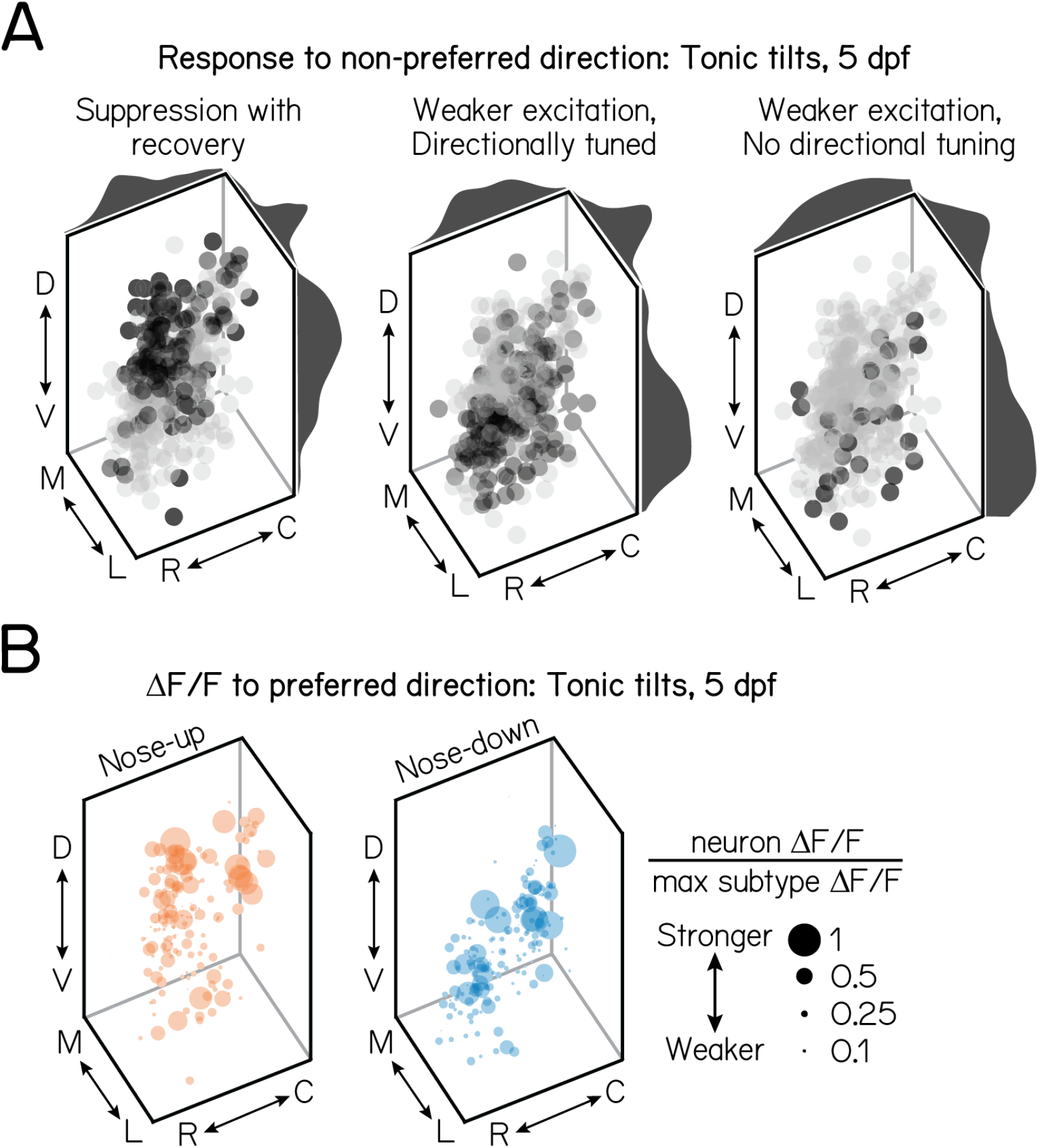
Projection neuron soma are locally organized according to similarities in tilt responses, Related to Figure 3. (A) Soma position of projection neurons with each response type to their non-preferred, tonic tilt direction. Dark grey shows neurons with the given response; light grey shows all other neurons. The directional tuning of neurons with weak excitation to non-preferred directions is determined by their directional tuning index (>0.1 or ≤0.1). Marginal distributions illustrate the probability of soma position in each spatial axis. Data from the same neurons shown in Figure 3G. // (B) Soma position of nose-up (left) and nose-down (right) projection neurons, scaled according to the strength of their calcium response (ΔF/F) to tilts relative to the strongest response observed for that subtype. Larger circles indicate stronger responses. Data from the same neurons shown in Figure 3.

**Table S1:** Number of synapses between 28 VIII^th^ nerve afferents (crista: semicircular canals; macula: utricular otoliths) and 19 projection neurons, Related to Figure 5. “Tuning” is an estimate of the sensitivity derived from the orientation of the macular hair cells, as described in ^55^; values between 0 and π/2 indicates rostral (nose-up) tuning. Projection neuron tuning (nose-up/nose-down) is defined as a weighted circular average of their utricular inputs. Projection neurons 1-3, 6, 11, 16-19 receive nose-up utricular input; neurons 6, 11, 16-19 also receive directionally-matched posterior cristae input. Projection neurons 4-5, 7-10, 12-15, 20 receive nose-down utricular input; neurons 5, 10, 13-15, and 20 also receive directionally-matched anterior cristae input.

| Afferent source | 1 | 2 | 3 | 4 | 5 | 6 | 7 | 8 | 9 | 10 | 11 | 12 | 13 | 14 | 15 | 16 | 17 | 18 | 19 | Tuning |
| --- | --- | --- | --- | --- | --- | --- | --- | --- | --- | --- | --- | --- | --- | --- | --- | --- | --- | --- | --- | --- |
| Ear_AnteriorCrista_R_01 | 0 | 0 | 0 | 0 | 0 | 0 | 0 | 0 | 0 | 0 | 0 | 0 | 8 | 0 | 0 | 0 | 0 | 0 | 0 |  |
| Ear_AnteriorCrista_R_02 | 0 | 0 | 0 | 0 | **2** | 0 | 0 | 0 | 0 | 0 | 0 | 0 | 0 | **8** | 0 | 0 | 0 | 0 | 0 |  |
| Ear_AnteriorCrista_R_03 | 0 | 0 | 0 | 0 | 0 | 0 | 0 | 0 | **3** | 0 | 0 | 0 | 0 | 0 | 0 | 0 | 0 | 0 | 0 |  |
| Ear_AnteriorCrista_R_04 | 0 | 0 | 0 | 0 | **1** | 0 | 0 | 0 | 0 | 0 | 0 | **9** | 0 | 0 | 0 | 0 | 0 | 0 | 0 |  |
| Ear_AnteriorCrista_R_05 | 0 | 0 | 0 | 0 | 0 | 0 | 0 | 0 | **2** | 0 | 0 | 0 | 0 | 0 | 0 | 0 | 0 | 0 | **12** |  |
| Ear_AnteriorCrista_R_06 | 0 | 0 | 0 | 0 | 0 | 0 | 0 | 0 | **5** | 0 | 0 | 0 | 0 | 0 | 0 | 0 | 0 | 0 | 0 |  |
| Ear_PosteriorCrista_R_01 | 0 | 0 | 0 | 0 | 0 | 0 | 0 | 0 | 0 | 0 | 0 | 0 | 0 | 0 | 0 | 0 | 0 | 0 | 0 |  |
| Ear_PosteriorCrista_R_02 | 0 | 0 | 0 | 0 | 0 | **2** | 0 | 0 | 0 | **2** | 0 | 0 | 0 | 0 | 0 | 0 | **8** | 0 | 0 |  |
| Ear_PosteriorCrista_R_03 | 0 | 0 | 0 | 0 | 0 | 0 | 0 | 0 | 0 | 0 | 0 | 0 | 0 | 0 | 0 | 0 | 0 | **7** | 0 |  |
| Ear_PosteriorCrista_R_04 | 0 | 0 | 0 | 0 | 0 | **3** | 0 | 0 | 0 | **1** | 0 | 0 | 0 | 0 | **10** | 0 | 0 | 0 | 0 |  |
| Ear_PosteriorCrista_R_05 | 0 | 0 | 0 | 0 | 0 | 0 | 0 | 0 | 0 | **1** | 0 | 0 | 0 | 0 | 0 | **5** | 0 | 0 | 0 |  |
| Ear_AnteriorMacula_R_01 | **1** | 0 | **2** | **3** | 0 | 0 | 0 | 0 | 0 | 0 | 0 | 0 | 0 | 0 | 0 | 0 | 0 | 0 | 0 | 6.00894 |
| Ear_AnteriorMacula_R_02 | 0 | **3** | 0 | 0 | 0 | 0 | 0 | 0 | 0 | 0 | 0 | 0 | 0 | 0 | 0 | 0 | 0 | 0 | 0 | 6.09989 |
| Ear_AnteriorMacula_R_03 | **1** | 0 | 0 | 0 | 0 | 0 | 0 | 0 | 0 | 0 | 0 | 0 | 0 | 0 | 0 | 0 | 0 | 0 | 0 | 6.08107 |
| Ear_AnteriorMacula_R_04 | 0 | **3** | **1** | 0 | 0 | **1** | 0 | 0 | 0 | 0 | 0 | 0 | 0 | 0 | 0 | 0 | 0 | 0 | 0 | 5.92756 |
| Ear_AnteriorMacula_R_05 | 0 | 0 | 0 | 0 | 0 | 0 | **1** | 0 | 0 | 0 | 0 | 0 | 0 | 0 | 0 | 0 | 0 | 0 | 0 | 0.204992 |
| Ear_AnteriorMacula_R_06 | **1** | 0 | 0 | 0 | **1** | 0 | **1** | 0 | 0 | 0 | 0 | 0 | 0 | 0 | 0 | 0 | 0 | 0 | 0 | 0.523691 |
| Ear_AnteriorMacula_R_07 | 0 | 0 | 0 | **1** | 0 | 0 | 0 | 0 | 0 | 0 | 0 | 0 | 0 | 0 | 0 | 0 | 0 | 0 | 0 | 0.825107 |
| Ear_AnteriorMacula_R_08 | 0 | 0 | 0 | 0 | **1** | 0 | 0 | 0 | 0 | 0 | **1** | 0 | 0 | 0 | 0 | 0 | 0 | 0 | 0 | 1.37515 |
| Ear_AnteriorMacula_R_09 | **1** | 0 | **4** | 0 | 0 | 0 | 0 | 0 | 0 | 0 | 0 | 0 | 0 | 0 | 0 | 0 | 0 | 0 | 0 | 6.04532 |
| Ear_AnteriorMacula_R_10 | 0 | 0 | 0 | 0 | 0 | 0 | 0 | 0 | 0 | 0 | 0 | 0 | 0 | 0 | 0 | 0 | 0 | 0 | 0 | 1.21203 |
| Ear_AnteriorMacula_R_11 | 0 | 0 | 0 | 0 | 0 | 0 | 0 | 0 | 0 | 0 | 0 | 0 | 0 | 0 | 0 | 0 | 0 | 0 | 0 | 4.20026 |
| Ear_AnteriorMacula_R_12 | 0 | 0 | 0 | **2** | 0 | 0 | 0 | **2** | 0 | 0 | 0 | 0 | 0 | 0 | 0 | 0 | 0 | 0 | 0 | 1.61526 |
| Ear_AnteriorMacula_R_13 | 0 | 0 | 0 | 0 | 0 | 0 | **1** | 0 | 0 | 0 | 0 | 0 | 0 | 0 | 0 | 0 | 0 | 0 | 0 | 0.257178 |
| Ear_AnteriorMacula_R_14 | 0 | 0 | 0 | 0 | 0 | 0 | 0 | 0 | 0 | 0 | 0 | 0 | 0 | 0 | 0 | 0 | 0 | 0 | 0 | 3.09177 |
| Ear_AnteriorMacula_R_15 | 0 | 0 | 0 | 0 | 0 | 0 | 0 | **2** | 0 | 0 | 0 | 0 | 0 | 0 | 0 | 0 | 0 | 0 | 0 | 0.93221 |
| Ear_AnteriorMacula_R_16 | 0 | 0 | 0 | 0 | 0 | 0 | 0 | 0 | 0 | 0 | 0 | 0 | 0 | 0 | 0 | 0 | 0 | 0 | 0 | 4.19152 |
| Ear_AnteriorMacula_R_17 | 0 | 0 | 0 | 0 | 0 | 0 | **1** | 0 | 0 | 0 | 0 | 0 | 0 | 0 | 0 | 0 | 0 | 0 | 0 | 1.31298 |

